# Bile Acid Scaffold Engineering Reveals an Androstane-Triol Derivative as a Potent Immunomodulator with Therapeutic Efficacy in EAE

**DOI:** 10.64898/2026.01.05.697637

**Authors:** Laura Calvo-Barreiro, Imane Boutitah-Benyaich, Herena Eixarch, Carmen Espejo, Moustafa T. Gabr

**Author notes:** Corresponding authors: Moustafa T. Gabr, e-mail address; Carmen Espejo.

## Abstract

**Background:** Neuroinflammation driven by dysregulated adaptive and innate immune responses plays a central role in the pathogenesis of multiple sclerosis and related autoimmune disorders of the central nervous system. While bile acids are increasingly recognized as endogenous immunomodulators, their therapeutic exploitation has been limited by modest potency and incomplete mechanistic understanding. Here, we report the rational engineering of a bile acid-derived scaffold that yields a potent small molecule immunomodulator with therapeutic efficacy in experimental autoimmune encephalomyelitis, a preclinical model of multiple sclerosis.

**Methods:** A focused series of bile acid-based compounds was established, leading to the identification of an androstane-triol derivative, BA59. The immunomodulatory activity of BA59 was evaluated using in vitro T cell differentiation assays, antigen-presenting cell phenotyping, and ex vivo immune profiling. Therapeutic efficacy was assessed in mice with established experimental autoimmune encephalomyelitis. Flow cytometry was used to characterize peripheral and central nervous system immune populations, including effector T cells, regulatory T cells, and antigen-presenting cells. Disease progression was monitored using clinical scoring and cumulative disease burden analyses.

**Results:** BA59 treatment significantly attenuated disease severity and cumulative disease burden when administered therapeutically after disease onset.

Immunophenotyping revealed a reduction in pro-inflammatory T helper 17 cells accompanied by an increase in regulatory T cells expressing the ectonucleotidase CD39. BA59 also reprogrammed antigen-presenting cells toward a tolerogenic phenotype, characterized by enhanced programmed death-ligand 1 expression. These immune changes were observed both in peripheral lymphoid tissues and within the central nervous system. Importantly, BA59 did not induce broad immunosuppression but instead reshaped immune checkpoint signaling and regulatory pathways associated with neuroinflammatory resolution.

**Conclusions:** This study identifies BA59 as a first-in-class androstane-triol immunomodulator that ameliorates experimental autoimmune encephalomyelitis through coordinated regulation of T cell balance, immune checkpoints, and antigen-presenting cell function. Our findings highlight bile acid scaffold engineering as a viable strategy for developing small molecule therapeutics that reprogram neuroinflammatory immune circuits, offering a promising translational approach for multiple sclerosis and related neuroinflammatory diseases.

## 1. Introduction

Multiple sclerosis (MS) is a chronic immune-mediated inflammatory disease of the central nervous system (CNS) characterized by demyelination and neurodegeneration, ultimately leading to disability progression [1, 2]. MS is a complex disease influenced by both genetic and environmental factors [2, 3]. While certain human leukocyte antigen (HLA) and non-HLA genes contribute to susceptibility [4–7], environmental factors such as viral infections, smoking, and vitamin D deficiency also play critical roles [8, 9]. Emerging evidence over the past two decades suggests that gut microbiota and its metabolites may be involved in MS pathogenesis [10–14], underscoring the need to better understand the immunological and metabolic pathways driving neuroinflammation. Although current therapies are partially effective at modulating immune activity and clinical relapses, they still lack significant impact on neurodegeneration, a key driver of progression in multiple sclerosis [2].

Bile acids (BAs) are cholesterol-derived metabolites synthesized in the liver and further transformed by the gut microbiota into diverse secondary BAs [15]. Beyond their classical roles in lipid absorption and enterohepatic circulation, secondary BAs have recently gained attention for their immunomodulatory properties influencing both innate and adaptive immune responses [16, 17]. Mechanistic studies have revealed that secondary BAs modulate adaptive immunity, particularly T helper (Th) 17 and regulatory T (Treg) cell differentiation, key drivers of autoimmune neuroinflammation. Specifically, lithocholic acid (LCA) derivatives such as dehydroLCA directly bind and inhibit the Th17 transcription factor RORγt, suppressing IL-17 production, while other derivatives including isoalloLCA promote Treg cell differentiation through mitochondrial metabolic signaling [18]. Additional bacterial-derived bile acid metabolite 3-epideoxycholic acid, an epimer of deoxycholic acid (DCA), enhances peripheral Treg cell induction via dendritic cell (DC) signaling and farnesoid X receptor (FXR)-dependent pathways [19]. *In vivo*, colonization with BA-transforming bacterial communities increases the proportion of Treg cell populations in the intestine and supports mucosal immune homeostasis [19].

Collectively, these findings position gut-derived BAs as key modulators of the Th17/Treg cell balance and suggest their potential to suppress inflammatory T cell responses in autoimmune diseases.

Recent studies have increasingly underscored the role of BA metabolism in modulating neuroinflammation and influencing disease progression in MS. Alterations in circulating levels of both primary and secondary BAs have been observed in adult and pediatric MS patients, with particularly pronounced reductions in individuals with progressive MS [20, 21]. Complementary findings from preclinical studies using the experimental autoimmune encephalomyelitis (EAE) model, a widely accepted animal model for MS, demonstrate that supplementation with secondary BAs such as tauroursodeoxycholic acid (TUDCA), DCA, and LCA can attenuate disease severity and promote regulatory immune responses [20, 22]. Moreover, clinical trials in progressive MS have demonstrated that TUDCA is safe and biologically active, modulating peripheral immune profiles and gut microbiota composition, and correlating with reduced CNS and retinal atrophy [21]. These findings underscore the therapeutic potential of BA derivatives in MS and support further exploration of their immunomodulatory mechanisms.

Previous studies have primarily explored naturally occurring BA metabolites for their immunomodulatory properties. In contrast, our work introduces a paradigm shift by applying rational scaffold engineering to design and synthesize a focused library of BA derivatives, enabling precise modulation of the immune pathways beyond the capabilities of native metabolites. We generated a of 64-BA derivative library and screened them *in vitro* under Th17- and Treg-polarizing conditions to identify potent immunomodulatory candidates. Two lead compounds were selected for *in vivo* evaluation in the EAE model, individually and in combination, to assess their effects on disease progression, CNS pathology, and immune phenotypes. Our findings identify an androstane-triol derivative with therapeutic potential in autoimmune neuroinflammation and uncover how BA-based scaffolds can modulate effector and Treg cell plasticity, antigen-presenting cell (APC) function, and immune checkpoint pathways. Ultimately, our goal is to identify BA-derived metabolites capable of reshaping T cell differentiation in ways that enhance or complement existing monoclonal therapies, targeting immune checkpoints or other key immunoregulatory receptors, in multiple sclerosis and other immune-mediated diseases.

## 2. Materials and Methods

### 2.1. Screened Library

A set of 64 compounds derived from secondary BAs isoalloLCA and dehydroLCA (also known as 3-oxoLCA) were selected from different vendors as suitable candidate library for the study of *in vitro* Th17 and Treg cell differentiation. All screened compounds were acquired as lyophilized powders, dissolved in 100% DMSO at a concentration equal to 5 or 10 mM and stored at -80°C until use (*Supporting Information, Table S1*).

For *in vivo* experiments, new lyophilized material of BA26 (Sarsasapogenin; Cat. # 24973, Cayman Chemical, Ann Arbor, MI, USA) and BA59 (Androstan-17-one, 3,6,7-trihydroxy-, (3β,5α,6α,7β)-; Cat. # AB03765, A2Bchem, San Diego, CA, USA) was reconstituted in DMSO at 16.13 mg/mL and stored at -20°C until use.

### 2.2. Mice

For the *in vitro* experiments, male and female C57BL/6J (Stock #: 000664), aged 5-6 weeks, were obtained from Jackson Laboratory (Farmington, CT, USA). Mice were housed under standard light- and climate-controlled conditions, and standard chow and water were provided *ad libitum*. All mouse experiments complied with ARRIVE (Animal Research: Reporting of In Vivo Experiments) guidelines and were carried out in accordance with the NIH (National Research Council) Guide for the Care and Use of Laboratory Animals and all procedures approved by the Institutional Animal Care and Use Committee (IACUC) at Weill Cornell Medicine (protocol number: 2022-0040).

For the *in vivo* experiments, C57BL/6JOlaHsd 8-week-old female mice purchased from Envigo (Venray, The Netherlands) were used. Mice were housed under standard light- and climate-controlled conditions, and standard chow and water were provided *ad libitum*. All experiments were performed in strict accordance with European Union (Directive 2010/63/EU) and Spanish regulations (Real Decreto 53/2013; Generalitat de Catalunya Decret 214/97). The Ethics Committee on Animal Experimentation of the Vall d’Hebron Research Institute approved all procedures described in this study (protocol number: CEEA 46/19: CEA-OH/9459R1/2).

### 2.3. *In Vitro* Differentiation of Mouse Naïve CD4⁺ **T Cells into Th17 and Treg Cell Subsets**

Spleens and lymph nodes were harvested from each C57BL/6J mouse and stored in ice-cold RPMI 1640 overnight at 4°C. 96-well flat-bottom plates were coated with 50 µL of 20 µg/mL goat anti-hamster IgG (Cat. # 0856984, MP Biomedicals, Santa Ana, CA, USA) and incubated overnight at 37°C. All procedures following mouse euthanasia and tissue harvest were conducted under sterile conditions in a tissue culture hood to prevent contamination. Sterile materials were used throughout.

For tissue processing, spleens and lymph nodes were dissociated mechanically using a syringe plunger in 3 mL of T cell media placed in a 70 µm cell strainer over a 60 mm Petri dish. T cell media was composed of XVIVO15 (Cat. # BW04-418Q, Lonza, Basel, Switzerland) at 96.35%, supplemented with HEPES (Cat. # 15630080, Gibco, Thermo Fisher, Waltham, MA, USA) at 1.00%, GlutaMAX (Cat. # 35-050-061, Gibco) at 1.00%, MEM Non-Essential Amino Acids Solution (Cat. # 11140050, Gibco) at 1.00%, Penicillin-Streptomycin (Cat. # P4333, Sigma-Aldrich, St. Louis, MO, USA) at 0.40%, and 1x PBS containing β-mercaptoethanol (Cat. # 21985023, Gibco) at 0.25%. The strainer was then transferred to a 50 mL conical tube, and the cell suspension was passed through. The Petri dish was washed with an additional 3 mL of T cell media, and the volume was added to the same tube. Subsequently, 1x PBS with 10% FBS was added up to 20 mL through the strainer, which was then discarded. Cells were centrifuged at 1,500 rpm for 5 minutes at 4°C. The pellet was resuspended in 1 mL of distilled water, followed by the addition of 1x PBS with 10% FBS up to 20 mL. The suspension was passed through a new 70 µm strainer into a fresh 50 mL conical tube and centrifuged again under the same conditions. The final pellet was resuspended in 500 µL of 1x PBS with 2% FBS, and cells were counted.

Naïve CD4⁺ T cells were isolated using the manual EasySep™ protocol according to the manufacturer’s instructions (EasySep™ Mouse Naïve CD4+ T Cell Isolation Kit, Cat. # 19765, StemCell, Vancouver, BC, Canada). Prior to isolation, cells were concentrated to 1×10⁸ cells/mL. After isolation, cells were counted and diluted in T cell media to a final concentration of 5×10⁵ cells/mL. For differentiation, 100 µL of this cell suspension (50,000 cells) was added per well. Each well received a total of 200 µL, consisting of 100 µL of cell suspension and 100 µL of 2x differentiation cocktail.

Experiments included two replicates per treatment and differentiation condition, with additional replicates for flow cytometry FMO (fluorescence minus one) controls. Treatments included a negative control, vehicle control (DMSO), and 64 test compounds derived from the secondary BAs isoalloLCA and dehydroLCA, as well as the two original BAs. Differentiation protocols included Th0 (control), Treg, and Th17 cell conditions.

Differentiation cocktails were prepared as follows: Th0 cell condition included 0.25 µg/mL anti-mouse CD3ε (Cat. # 100302, BioLegend, San Diego, CA, USA), 1 µg/mL anti-mouse CD28 (Cat. # 102102, BioLegend), and 20 U/mL recombinant human IL-2 (Cat. # 200-02, PeproTech, Thermo Fisher). Th17 cell condition included 0.25 µg/mL anti-mouse CD3ε, 1 µg/mL anti-mouse CD28, 10 ng/mL recombinant mouse IL-6 (Cat. # 216-16, PeproTech), 0.5 ng/mL recombinant human TGF-β1 (Cat. # 100-21, PeproTech), 1 µg/mL anti-mouse IL-4 (Cat. # 504122, BioLegend), and 1 µg/mL anti-mouse IFN-γ (Cat. # 505834, BioLegend). Treg cell condition included 0.25 µg/mL anti-mouse CD3ε, 1 µg/mL anti-mouse CD28, 20 U/mL recombinant human IL-2, 0.02 ng/mL recombinant human TGF-β1, 1 µg/mL anti-mouse IL-4, and 1 µg/mL anti-mouse IFN-γ.

Prior to cell seeding, the coating solution was aspirated from the 96-well plate and wells were washed once with 1x PBS. Then, 100 µL of 2x differentiation cocktail was added to each well, followed by 100 µL of 2x cell suspension. Uniform cell density across wells was confirmed under brightfield microscopy. Plates were incubated at 37°C for 16 hours. After incubation, 10x bile acid solutions (200 μM, containing 4% DMSO) were prepared in T cell media, vortexed, and sonicated to ensure solubilization.

Subsequently, 20 μL of the freshly prepared 10x bile acid solutions were added to the appropriate wells, resulting in a final concentration of 20 μM and 0.4% DMSO. BA65 (dehydroLCA) and BA30 (isoalloLCA acid) were used as positive controls for Th17 and Treg cell differentiation, respectively. Plates were returned to the incubator for an additional two days.

After 72h *in vitro*, cells requiring activation were transferred to 96-well round-bottom plates. Cells were gently detached by pipetting up and down 10-20 times, centrifuged at 450xg for 5 minutes, and the supernatant was discarded. Pellets were resuspended in 200 µL of 1x T cell stimulation media [50 ng/mL phorbol 12-myristate 13-acetate (PMA, Cat. # P1585, Sigma-Aldrich), 1 µM ionomycin (Cat. # I0634, Sigma-Aldrich), 1 µL/mL of GolgiPlug (Cat. # 555029, BD Biosciences, Franklin Lakes, NJ, USA) and 0.67 µL/mL GolgiStop (Cat. # 554724, BD Biosciences)] and incubated for 4 hours at 37°C. Following stimulation, cells were stained as needed and analyzed by flow cytometry.

### 2.4. Induction and Assessment of EAE

Anaesthetized mice were immunized by subcutaneous injection of 100 μL of PBS containing 100 μg of MOG_35-55_ (Proteomics Section, Universitat Pompeu Fabra, Barcelona, Spain) emulsified in 100 μL of complete Freund’s adjuvant [incomplete Freund’s adjuvant (IFA, Sigma-Aldrich) containing 4 mg/ml *Mycobacterium tuberculosis* H37RA (BD Biosciences)]. At 0- and 2-day postimmunization (dpi), mice were intravenously injected with 250 ng of *pertussis* toxin (Sigma-Aldrich).

Mice were weighed and examined daily for neurological signs in a blinded manner using the following criteria: 0, no clinical signs; 0.5, partial loss of the tail tonus for 2 consecutive days; 1, paralysis of the whole tail; 2, mild paraparesis of one or both hind limbs; 2.5, severe paraparesis or paraplegia; 3, mild tetraparesis; 4, severe tetraparesis (paraplegia in hind limbs); 4.5, severe tetraparesis (incapable of turning around); 5, tetraplegia; and 6, death.

Corrective measures and endpoint criteria to ensure the welfare of EAE-diseased animals included: i) wet food pellets in the bed-cage to facilitate access to food as well as hydration, ii) subcutaneous administration of 0.5 ml of glucosaline serum (glucose 10%) in case of more than 15% of weight loss, and iii) mouse euthanasia if the weight loss exceeded 30% or an animal reached the clinical score of 5.

The overall clinical score per mouse was calculated as the area under the curve (AUC) of the daily clinical score throughout the experiment.

All data presented are in accordance with the guidelines suggested for EAE publications [23] and the ARRIVE (*Animal Research: Reporting of In Vivo Experiments*) guidelines for animal research [24, 25].

### 2.5. EAE Experimental Design and Compound Administration

Before therapeutic treatment, mice were randomized into clinically equivalent groups upon reaching a clinical score equal to or greater than 1, between 12- and 19-days post-immunization (dpi). Sarsasapogenin (BA26) and androstan-17-one, 3,6,7-trihydroxy-, (3β,5α,6α,7β)- (BA59) were administered orally once daily via gavage at 5 mg/kg or 25 mg/kg in 200 μL of either compound solution or vehicle (DMSO), until the end of the experiment (28 dpi).

### 2.6. Evaluation of Motor Function via Rotarod Testing

At 28 dpi, motor performance was assessed using a Rotarod apparatus (Ugo Basile, Gemonio, Italy). The device was programmed to accelerate from 4 to 40 rpm over a 300-s trial. Mice were placed on the rotating cylinder, and the latency to fall was recorded for each trial. For every animal, motor performance was expressed as the mean latency to fall across three consecutive trials.

### 2.7. Flow Cytometry

In the *in vitro* Th17/Treg cell differentiation experiments, antibody-labelled T cells were acquired in a Fortessa flow cytometer (Becton-Dickinson, Franklin Lakes, NJ, USA) and the analysis was performed using the FlowJo™ v10.10 Software (BD). Cell subsets were analyzed using fluorochrome-conjugated monoclonal antibodies (mAbs) after discrimination of dead cells by Fixable Viability Stain (BD Pharmingen, BD Biosciences). *Ex vivo* stimulation of splenocytes was performed with 50 ng/ml PMA (Sigma-Aldrich) and 1 μM ionomycin (Sigma-Aldrich) in the presence of GolgiPlug (BD Biosciences) and GolgiStop (BD Biosciences) for 6 h. Then, CD3ε (Cat. # 553061, BD Pharmingen), CD4 (Cat. # 46-0042, Invitrogen, Thermo Fisher), and CD8a (Cat. # 560182, BD Pharmingen) staining was performed. For analysis of the Treg cell population, CD25 (Cat. # 12-0251, Invitrogen) and FOXP3 (Cat. # 17-5773, Invitrogen) intracellular staining were additionally performed. For Th17 analysis, cytokine intracellular staining was performed using fluorochrome-labelled anti-IL-17A (Cat. # 512322, Biolegend) mAb.

In the EAE experiments, spleen cell suspensions were prepared at the end of the experiment (28 dpi) as described previously for *in vitro* experiments. Antibody-labelled splenocytes were acquired in a Cytoflex flow cytometer (Beckman Coulter, Brea, CA, USA) and the analysis was performed using the FlowJo™ v10.10 Software (BD). Cell subsets were analyzed using fluorochrome-conjugated mAbs after discrimination of dead cells by Fixable Viability Stain (BD Pharmingen). For analysis of the Treg cell population, CD3ε (Cat. # 553061, BD Pharmingen), CD4 (Cat. # 560181, BD Pharmingen), CD8 (Cat. # 563152, BD Horizon) and CD25 (Cat. # 12-0251, eBioscience, San Diego, CA, USA) were used. FoxP3 intracellular staining was performed using fluorochrome-labelled anti-FoxP3 mAb (Cat. # 17-5773, eBioscience). Additionally, CD39 (Cat. # 25-0391, eBioscience) was also selected and evaluated in the Treg cell population. For DC subpopulations and activation status, mAbs specific for B220 (Cat. # 553088, BD Pharmingen), CD11b (Cat. # 25-0112-82, eBiosciences), CD11c (Cat. # 47-0114-82, eBiosciences), CD8a (Cat. # 563152, BD Horizon), CD80 (Cat. # 553769, BD Pharmingen), CD86 (Cat. # 563077, BD Horizon), major histocompatibility complex class II (MHCII, Cat. # 562363, BD Pharmingen), and PD-L1 (CD274, Cat. # 124315, Biolegend) were used. For analysis of T cell subpopulations and immune checkpoints, CD3ε (Cat. # 553061, BD Pharmingen), CD4 (Cat. # 560181, BD Pharmingen), CD8a (Cat. # 563068, BD Horizon), CD44 (Cat. # 560570, BD Pharmingen), CD62L (Cat. # 563252, BD Horizon), CTLA-4 (CD152, Cat. # 564331, BD Pharmingen), LAG-3 (CD223, Cat. # 552380, BD Pharmingen), PD-1 (CD279, Cat. # 135217, Biolegend), and TIM-3 (CD366, Cat. # 25-5870, eBioscience) were selected and assessed. For intracellular cytokine determination, *ex vivo* stimulation of splenocytes was performed with 50 ng/ml PMA (Sigma-Aldrich) and 1 μg/ml ionomycin (Sigma-Aldrich) in the presence of GolgiPlug (BD Biosciences) and GolgiStop (BD Biosciences) for 6 h. Then, CD3ε (Cat. # 553061, BD Pharmingen), CD4 (Cat. # 46-0042, eBioscience), and CD8a (Cat. # 47-0081, eBioscience) staining was performed. Cytokine intracellular staining was performed using fluorochrome-labelled anti-IFN-γ (Cat. # 554413, BD Pharmingen), anti-TNF-α (Cat. # 506339, Biolegend), anti-IL-10 (Cat. # 563276, BD Horizon), and anti-IL-17A (Cat. # 559502, BD Pharmingen) mAbs and anti-CD69 mAb (Cat. # 563290, BD Horizon) to assess proper *ex vivo* stimulation.

The different immune cell populations were defined as the percentage of positive cells for a certain marker or combination of markers within the parent population. The level of expression of an inducible cell marker was defined as the median fluorescence intensity (MFI) for a certain cell marker within the cell marker-positive parent population.

### 2.8. Histopathological Analysis

At the end of the experiment (28 dpi), spinal cords of euthanized EAE mice were collected, fixed in a 4% paraformaldehyde solution, embedded in paraffin, and cut into 4-μm thick coronal sections.

Inflammatory infiltration was assessed by haematoxylin and eosin (H&E) staining. Inflammatory infiltration was scored as follows: 0, no lesion; 1, cellular infiltration only in the meninges; 2, very discrete and superficial infiltrates in parenchyma; 3, moderate infiltrate (less than 25%) in the white matter; 4, severe infiltrates (less than 50%) in the white matter; and 5, more severe infiltrates (more than 50%) in the white matter.

Demyelination was assessed by rabbit polyclonal anti-myelin basic protein (MBP, Cat. # AB7349, Merck, Kenilworth, NJ, USA). Briefly, coronal spinal cord sections were deparaffinized and rehydrated, and antigen retrieval and preincubation in blocking solution for 1 h at room temperature were performed. Immunofluorescences were carried out by incubating sections with the corresponding primary antibody diluted in the blocking solution overnight at 4 °C. After rinsing, sections were incubated with the goat anti-rabbit Alexa 568 antibody (Cat. # A11077, Invitrogen, Carlsbad, CA, USA) in the blocking solution for 1 h at room temperature. Finally, cell nuclei were stained with 4’,6-diamidino-2-phenylindole (DAPI, Cat. # D9542, Sigma-Aldrich) and coverslips were mounted with Fluoromount-G (Cat. # 00-4958-02, Invitrogen).

Images were acquired using a Thunder fluorescence microscope (Leica Microsystems, Wetzlar, Germany) and LAS X visualization software (Leica Microsystems), and mosaic images were obtained at a magnification of 20x and analyzed using SlideViewer and ImageJ 1.54p. For every staining condition, two mosaic images from the thoracic spinal cord, each separated by 300 μm, were selected per mouse and evaluated in a blinded manner. For inflammatory infiltration measurements, the results are shown as the individual score per mouse. For demyelination measurements, the results are shown as the percentage of white matter area without MBP staining relative to the total white matter area.

## 3. Results

### 3.1. Focused Screening of a 64-Member Bile Acid Library Reveals Novel Modulators of CD4⁺ T Cell Differentiation

Two secondary BAs, dehydroLCA and isoalloLCA, had been previously characterized for their immunomodulatory properties [18]. In prior studies, both compounds reduced Th17 cell differentiation, whereas isoalloLCA uniquely enhanced Treg cell differentiation. Notably, these functional differences arise from a minimal structural variation: a hydroxy group versus an oxo group at the C3 position of ring A. This subtle change, yet distinct qualitative activity (the hydroxy group modulating both Th17 and Treg cell pathways, the oxo group acting selectively on Th17 cells), suggested that even small modifications around the gonane core could produce significant shifts in CD4⁺ T cell differentiation.

Guided by this structure-function relationship, we designed a focused 64-member custom library (Table S1). All compounds retained the gonane steroid nucleus and a hydroxy or oxo substituent at C3 (R1), while introducing systematic variations in oxidation state, functional groups, and stereochemistry at positions R2-R6 (Figure 1A). This library was screened *in vitro* under Th17- and Treg-polarizing conditions, with Th0 cell cultures serving as controls. Naïve CD4⁺ T cells were first incubated for 16 h under the appropriate differentiating conditions. Later, compounds were added (20 μM) and cells were cultured for two additional days, for a total of 72 h before assessing differentiation outcomes (Figure 1B).

**Figure 1.**
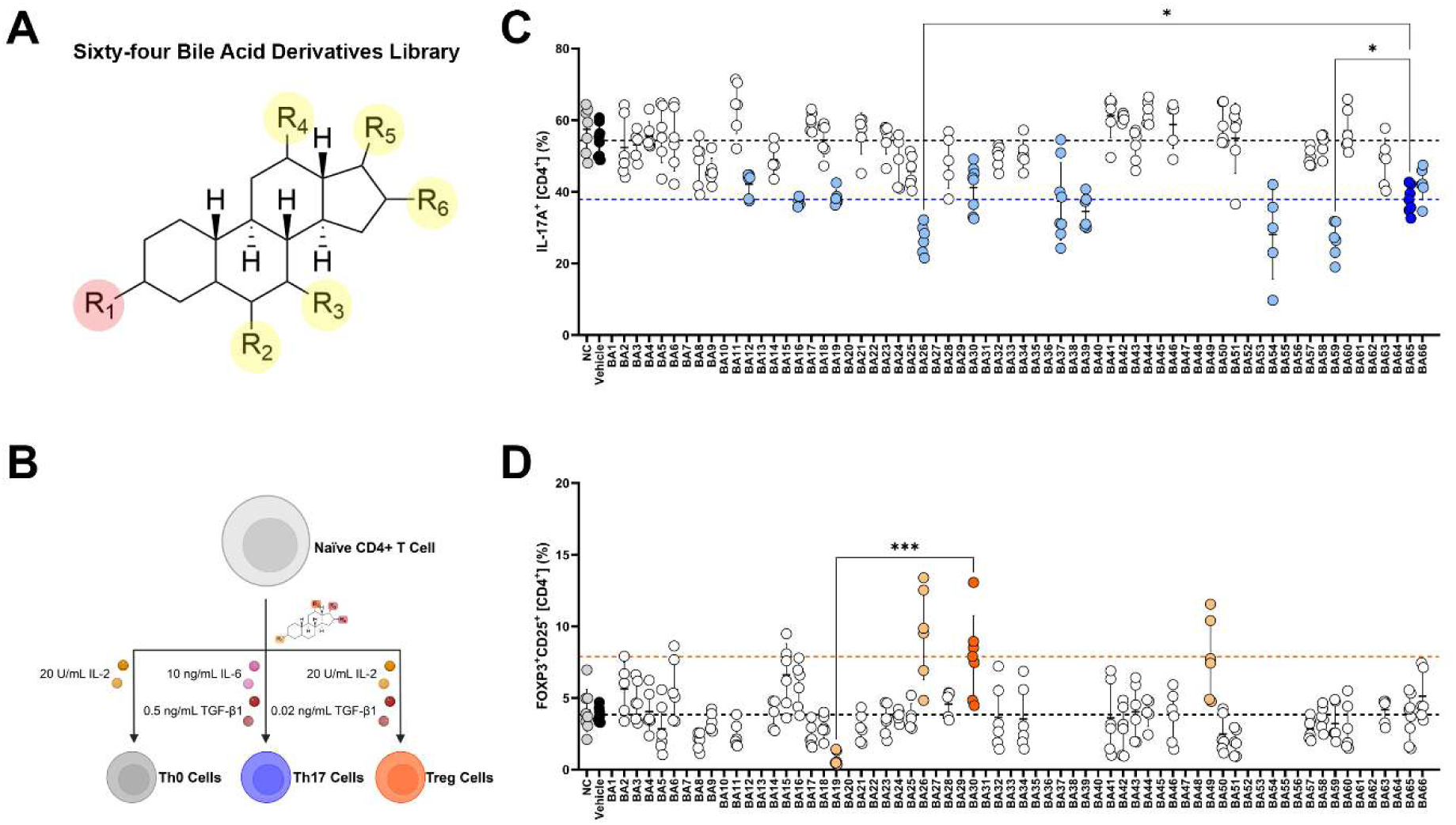
Novel BA derivatives match or exceed the effects of known secondary BAs on Th17 and Treg cell differentiation. (**A**) Schematic of the 64-member focused BA derivative library. All compounds maintain the gonane steroid nucleus and a hydroxy or oxo substituent at C3 (R1), with systematic variation in oxidation state, functional groups, and stereochemistry at positions R2-R6. (**B**) Workflow for *in vitro* CD4⁺ T-cell differentiation. Lymph nodes and spleens were harvested from naïve C57BL/6J mice, mechanically dissociated, and naïve CD4⁺ T cells were isolated by immunomagnetic negative selection. Cells were cultured under Th0, Th17, or Treg cell polarizing conditions. After 16 h, individual BA derivatives (20 μM; 0.4% DMSO) or vehicle (0.4% DMSO) were added and cultures were continued for two additional days (72 h total) before flow cytometry analysis. (**C**) Percentage of Th17 cells after 72 h under Th17-polarizing conditions. NC (negative control) represents differentiation in the absence of DMSO and vehicle denotes differentiation in 0.4% DMSO (matching BA conditions). Each BA was tested at 20 μM. BA65 (dehydroLCA, dark blue) served as the positive control. Compounds shown in light blue significantly differed from vehicle (*p value* < 0.05), and connecting brackets indicate compounds significantly different from BA65. (**D**) Percentage of Treg cells after 72 h under Treg-polarizing conditions. NC and vehicle controls are defined as in (C). BA30 (isoalloLCA, dark orange) served as the positive control. Compounds in light orange significantly differed from vehicle (*p value* < 0.05), and connecting brackets indicate compounds significantly different from BA30. The experimental data represents four independent experiments with six to eight replicates per experimental condition. Individual values for each replicate are displayed, along with the mean and standard deviation across all samples. Statistical analyses were performed using one-way ANOVA with Dunnett’s multiple comparison test. * *p value* < 0.05; *** *p value* < 0.001.

Before evaluating the effects of the library on Tncell differentiation, we assessed cell viability for each compound (Figures S1-S3). Compounds that reduced viability at 20 μM relative to the vehicle control were excluded from subsequent analyses; therefore, they do not appear in Figures 1C and 1D. As expected, the positive controls dehydroLCA and isoalloLCA reproduced their previously reported activities: both reduced Th17 cell differentiation, and isoalloLCA additionally enhanced Treg cell differentiation (Figures 1C, D). Among the library members, nine compounds decreased Th17 cell frequencies (light blue, Figure 1C), and three altered Treg cell differentiation (light orange, Figure 1D) compared with the vehicle.

For Th17 cell differentiation, two compounds, BA26 (sarsasapogenin) and BA59 (an androstane-triol derivative), showed greater activity than the positive control dehydroLCA (BA65) (Figure 1C). Relative to the vehicle condition (54.31 ± 4.52% Th17 cells, n=8), dehydroLCA reduced Th17 cell frequencies to 37.90 ± 3.80% (n=7), whereas BA26 and BA59 decreased them to 26.86 ± 4.06% (n=6) and 26.52 ± 4.95% (n=6), respectively. Thus, BA65 reduced Th17 cells by 30.23 ± 6.99%, while BA26 and BA59 achieved 50.54 ± 7.48% and 51.17 ± 9.11% reductions. When directly compared to BA65, BA26 and BA59 further decreased Th17 cell frequencies by 29.12 ± 10.72% and 30.01 ± 13.06%, respectively. Under Th0 cell conditions, Th17 cell frequencies remained below 1% of CD4⁺ T cells and were unaffected by any compound, including the positive control (Figure S4).

For Treg cell differentiation, isoalloLCA (BA30) induced an increase from 3.84 ± 0.53% (n=7) in the vehicle group to 7.88 ± 2.87% (n=7), corresponding to a 205.32 ± 74.70% increase (Figure 1D). BA26 and BA49 yielded Treg frequencies of 9.51 ± 3.25% (n=6) and 7.80 ± 2.78% (n=6), with relative increases of 247.66 ± 84.64% and 203.13 ± 72.31%, respectively. Although BA26 produced modest improvements over the positive control (20.62 ± 41.22%), that difference did not reach statistical significance. BA19, in contrast, markedly reduced Treg cell differentiation to 0.76 ± 0.48% (n=6), an 80.12 ± 12.46% decrease relative to the vehicle. Under Th0 cell conditions, Treg cell frequencies were low (0.39 ± 0.18%, n=7) for the vehicle condition but increased to 1.50 ± 0.76% (n=8) and 0.79 ± 0.18% (n=6) with BA26 and BA49, compared to 1.00 ± 0.35% (n=8) with BA30 (Figure S5). Despite the absence of TGF-β1 in the Th0-conditioned media, BA26 and BA49 increased Treg cell frequencies by 3.83-fold and 2.02-fold, respectively, and BA30 by 2.56-fold, mirroring the trends observed under Treg-polarizing conditions.

### 3.2. Dose-Response Characterization of Selected Bile Acid Derivatives in Th17 and Treg Cell Differentiation

Building upon the initial single-dose experiments, we next wanted to determine whether the BA derivatives exhibited a dose-dependent effect. Although several compounds showed clear activity at 20 μM, dose-response experiments (ranging from 1.28 to 50 μM) were performed to assess robustness, reproducibility, dose-dependency, and potency, as well as to distinguish specific immunomodulatory effects from nonspecific toxicity. These data also enable quantitative comparisons essential for structure-activity relationship (SAR) analyses and translational evaluation.

Regarding Th17 cell modulation, ten BAs (BA12, BA16, BA19, BA26, BA30, BA37, BA39, BA54, BA59, and BA66), along with the positive control BA65, demonstrated clear dose-response trends in their inhibition of Th17 cell differentiation (Figure S7). All compounds except BA16, BA59, and BA65 showed cytotoxicity at 50 μM (Figure S6). Given their superior activity in the single-dose screen, BA26 and BA59 were further evaluated and compared directly with the positive control (Figures 2A, C). BA26 consistently outperformed BA65 at 20 μM and displayed a dose-dependent effect. However, its potency could not be reliably calculated because only the four non-toxic concentrations could be included in the fit, preventing accurate determination of the lower plateau (Figure 2A). BA59 yielded a well-defined sigmoidal curve with an IC_50_ of 24.04 μM (Bottom = -70.12%, Top = -10.39%, Hill slope = -2.996, R^2^ = 0.92), outperforming the positive control BA65, whose IC_50_ was 34.24 μM (Bottom = -111.8%, Top = -15.87%, Hill slope = -2.096, R^2^ = 0.93) (Figure 2A).

**Figure 2.**
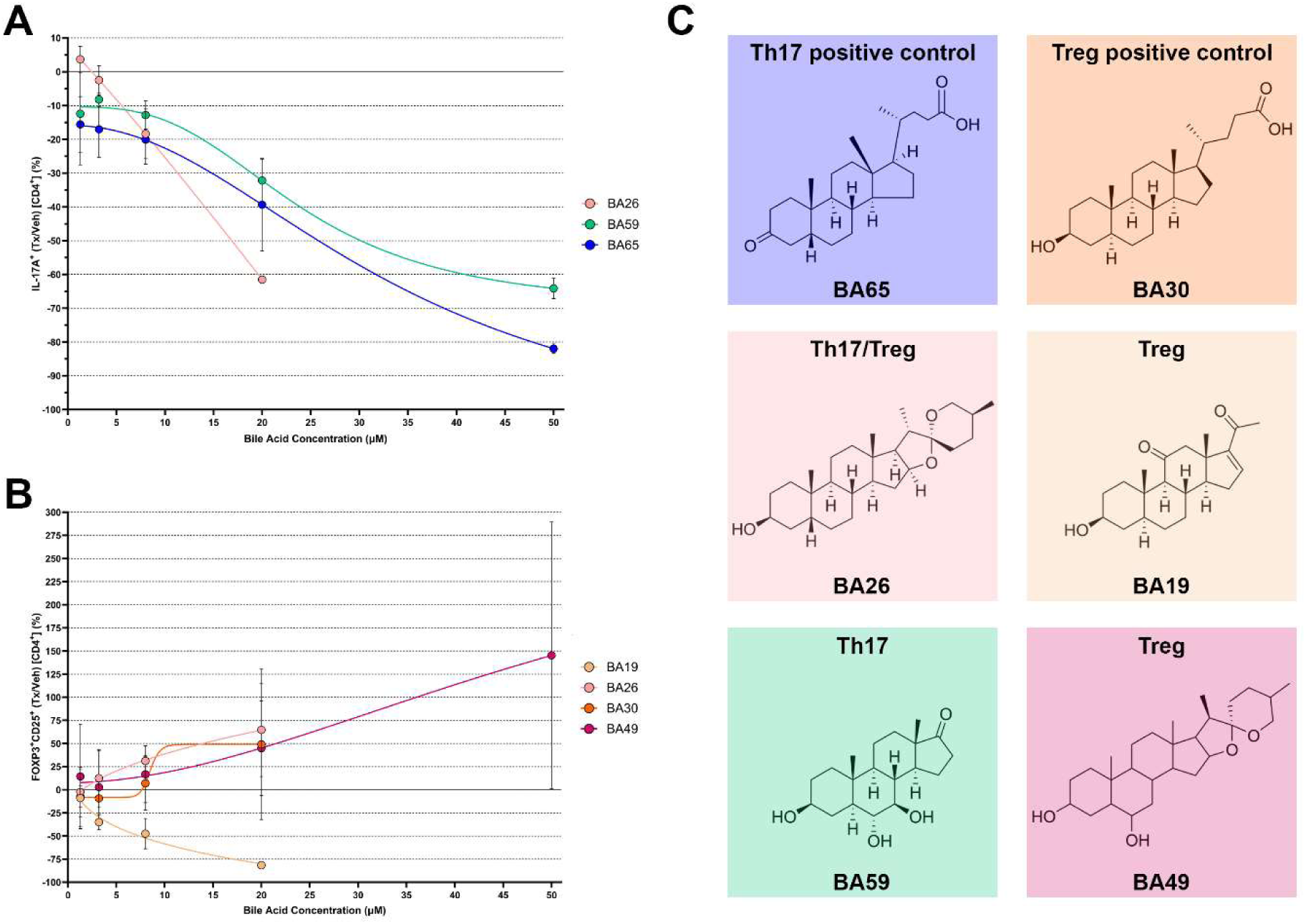
Selected BA derivatives exhibit dose-dependent effects on CD4^+^ T cell differentiation. Using the same experimental workflow as in Figure 1, naïve CD4^+^ T cells were cultured under Th17- or Treg-polarizing conditions and treated with increasing concentrations: 1.28 µM, 3.2 µM, 8 µM, 20 µM (same as in the single-dose screening), and 50 µM; of previously identified active BAs. (**A**) BA26 (sarsasapogenin) and BA59 (an androstane-triol derivative), as well as the positive control BA65 (dehydroLCA), reduced Th17 cell population frequency in a dose-dependent manner within the non-toxic range for each candidate compound. (**B**) BA26 (sarsasapogenin) and BA49 (a spirostan-3,6-diol derivative), as well as the positive control BA30 (isoalloLCA), increased Treg cells frequency in a dose-dependent manner within the non-toxic range for each candidate compound. On the other hand, BA19 (11-oxo-pregnenolone derivative) decreased Treg cell population in a dose-response manner within its non-toxic range. (**C**) Chemical structures of the main secondary BA derivatives (BA19, BA26, BA49, and BA59) and the positive controls (BA30 and BA65) that exert immunoregulatory effects on Th17 and/or Treg cell populations. Data represent three independent experiments, each with one technical replicate. Mean values and standard deviations for each concentration are shown. EC_50_/IC_50_ values were calculated by fitting dose-response data to a sigmoidal four-parameter logistic (4PL) model, using BA concentration as the X variable. EC_50_/IC_50_ values were derived from 5-point dose-response curves; when compounds exhibited reduced viability at the highest concentration, only the four non-toxic concentrations were included in the fit.

On the Treg cell side, three derivatives (BA19, BA26, and BA49) and the positive control BA30 were examined for effects on Treg cell differentiation (Figures 2B, C). All compounds exhibited dose-response behavior, and all except BA49 were cytotoxic at 50 μM (Figures 2B, S8). BA49 showed an EC_50_ of 57.55 μM (Bottom = 7.62%, Top = 325.2%, Hill slope = 1.902, R^2^ = 0.42). The positive control BA30 produced an apparent EC_50_ of 8.49 μM, but this value should be interpreted with caution: only four non-toxic concentrations could be included, the hill slope was unrealistically steep (17.05), and increased variability at 20 μM compromised model stability. Although the curve displayed a sigmoidal shape, the upper plateau was not clearly defined, making the EC_50_ only approximate. For BA19 and BA26, IC_50_/EC_50_ values could not be reliably calculated; however, BA19 induced a reproducible decrease in Treg cell frequency across the tested concentration range, while BA26 consistently induced Treg cell levels comparable to those of the positive control at 20 μM (Figure 2B).

Taken everything into account, these results led us to select BA59 as the leading candidate for Th17 cell reduction, and BA26 as a dual-acting compound capable of increasing Treg cells while decreasing Th17 cell populations. Based on their immunomodulatory properties, we considered these compounds particularly promising for translational evaluation in autoimmune disease models.

### 3.3. Oral Administration of BA59 and in Combination with BA26 Reduces Clinical Disease Severity in EAE

Based on the *in vitro* activity of our 64-member BA-derived library, we selected two candidates, BA26 and BA59, for *in vivo* testing in the EAE model (Figure 3A). BA26 was prioritized because it simultaneously enhanced Treg cell differentiation and reduced Th17 polarization *in vitro*, whereas BA59 was chosen for its robust and selective suppression of Th17 cell differentiation. To determine whether these immunomodulatory properties translated into therapeutic benefit, we first evaluated each compound individually in EAE. Mice were therapeutically treated daily beginning at a clinical score equal to or higher than 1 with 5 mg/kg BA26 or 25 mg/kg BA59. Given BA26’s dual *in vitro* properties relative to BA59, we opted to begin with a lower dose. Under these conditions, BA59 produced a measurable reduction in clinical severity after 10 days of treatment, leading to an overall trend toward clinical improvement (AUC = 24.29 ± 10.20, n = 12, *p* = 0.093) compared with vehicle-treated controls (33.08 ± 10.58, n = 12) (Figure S9). In contrast, BA26 did not show any detectable clinical benefit (32.29 ± 11.22, n = 12) (Figure S9).

**Figure 3.**
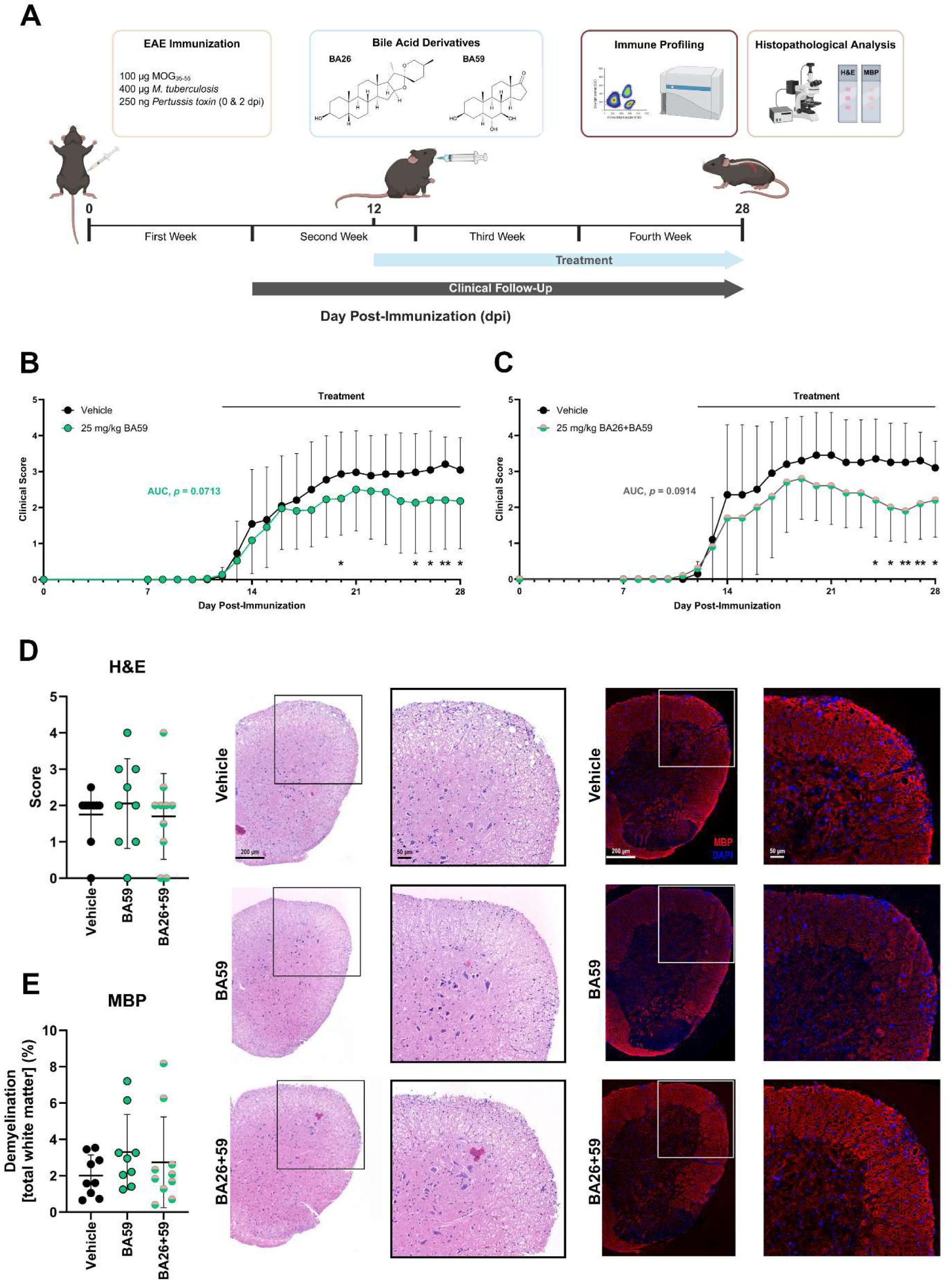
Oral treatment with BA59 and its combination with BA26 improve disease parameters in EAE mice. (**A**) Following EAE induction with MOG_35-55_, complete Freund’s adjuvant, and *pertussis* toxin, mice were randomized into clinically equivalent groups upon reaching a clinical score equal to or greater than 1 (12-19 dpi). BA26 (sarsasapogenin) and BA59 (an androstane-triol derivative) were administered once daily by oral gavage at 5 or 25 mg/kg in 200 μL of compound solution or vehicle (DMSO) until the end of the experiment (28 dpi). Treatments were given individually and in combination. Spleens and spinal cords were collected for immune profiling and histopathological analysis at the end of the experiment. (**B**) Therapeutic administration of BA59 (25 mg/kg) reduced disease severity from 25 dpi and decreased the AUC of clinical scores compared to vehicle (30.73 ± 15.27 vs. 38.95 ± 14.21; *p value* = 0.071). Data represents two independent experiments (n = 22 animals/treatment) and is shown as the mean ± SD. Statistical analyses were performed using Welch’s t-test. * *p value* < 0.05; ** *p value* < 0.01. (**C**) Combined treatment with BA26+BA59 (25 mg/kg each) reduced disease severity from 24 dpi and lowered the AUC of clinical scores compared to vehicle (33.80 ± 15.33 vs. 46.00 ± 15.26; *p value* = 0.091). Data represents a single experiment (n = 10 animals/treatment) and is shown as the mean ± SD. Statistical analyses were performed using Welch’s t-test. * *p value* < 0.05; ** *p value* < 0.01. (**D**) Overall inflammation assessed by H&E staining showed no differences among treatment groups at 28 dpi. The graph shows the mean inflammatory score per animal (two coronal sections per animal) from one experiment (Vehicle, n=10; and BA59, n=9; BA26+59, n=10). Data are also presented as mean ± SD for each experimental group. (**E**) Demyelination assessed by MBP staining showed no differences among experimental groups at 28 dpi. The graph shows the mean percentage of demyelination relative to total white matter area (two coronal sections per animal) from the same experiment (Vehicle, n=9; and BA59, n=9; BA26+59, n=10). Data are also presented as mean ± SD for each experimental group.

To assess whether insufficient exposure accounted for the lack of efficacy of BA26, we performed a second experiment in which both BA26 and BA59 were administered at 25 mg/kg. Consistent with our initial findings, BA59 again significantly ameliorated experimental disease and showed a clear trend toward overall clinical improvement (AUC = 30.73 ± 15.27, n = 22 vs. 38.95 ± 14.21, n = 22, *p* = 0.0713)

(Figure 3B), whereas BA26 remained ineffective even at the higher dose (Figure S10). Because BA26 showed supplementary immunomodulatory effects compared to BA59, we also tested the compounds in combination, administering both BA derivatives at 25 mg/kg beginning once clinical signs were already established (clinical score ≥ 1).

Notably, co-treatment recapitulated the beneficial effect observed with BA59 alone, resulting in a significant reduction in disease severity and a trend toward overall clinical improvement compared with vehicle controls (AUC = 33.80 ± 15.33, n = 10 vs. 46.00 ± 15.26, n = 10, *p* = 0.0914) (Figure 3C). Rotarod assessment revealed no differences in motor function between treatment groups at the end of the experiment (28 dpi), neither BA59 alone nor in combination with BA26 differed from the vehicle-treated mice (Figure S11).

Together, these results indicate that BA59 exhibits reproducible *in vivo* efficacy in EAE, consistent with its potent anti-Th17 activity *in vitro*, while BA26 alone is insufficient to confer clinical benefit but does not interfere with BA59-mediated clinical improvement when co-administered. Finally, histopathological analysis of inflammation and demyelination of the spinal cords at the end of the experiment did not show statistically significant differences between groups in overall inflammatory infiltrates or white matter demyelination (Figures 3D, E).

### 3.4. BA59 Induces Broad Modulation of Peripheral Immune Cell Populations in EAE

To investigate how BA treatment modulates immune responses associated with the observed clinical improvement in EAE, we performed multiparameter flow cytometry analysis of major immune cell compartments, including: CD4⁺ and CD8⁺ T cells, B cells, and DCs; at the chronic phase of disease (28 dpi) (Figures S12-S16). At this time point, mice had already received oral treatment for more than 14 days and exhibited sustained clinical benefit, enabling assessment of immunological correlates underlying therapeutic efficacy.

To determine how BA treatment modulated effector and Treg cell responses, we quantified cytokine-expressing CD4⁺ and CD8⁺ T cell subsets. BA59 monotherapy markedly shifted the CD4⁺ T cell compartment toward a less inflammatory phenotype, as shown by a significant reduction in the IFN-γ⁺TNF-α⁻/IL-10⁺ ratio and increased CD4/CD8 ratios within both IFN-γ⁺TNF-α⁺ and IFN-γ⁺TNF-α⁻ populations (Figures 4A-C). These changes indicate a preferential modulation of CD4⁺ effector responses relative to CD8⁺ cells. BA59 also expanded IFN-γ⁺TNF-α⁻ CD8⁺ T cells, a partially activated effector subset associated with attenuated cytotoxicity, further supporting a shift toward a less inflammatory effector profile (Figure 4D). In contrast, the combined treatment failed to reproduce the immunomodulatory effects of BA59 alone. The combination neither decreased the IFN-γ⁺TNF-α⁻/IL-10⁺ CD4⁺ ratio nor increased CD4-biased effector ratios (Figures 4A-C). Instead, it reduced the IFN-γ⁻TNF-α⁺ CD4/CD8 ratio while decreasing IFN-γ⁺TNF-α⁻ CD8⁺ cells, suggesting a shift toward CD8-driven TNF-α⁺ responses (Figures 4D, E). Together, these data demonstrate that BA59 selectively reprograms both CD4⁺ and CD8⁺ T cell cytokine profiles toward a less inflammatory state, whereas co-administration with BA26 counteracts these beneficial immunological effects.

**Figure 4.**
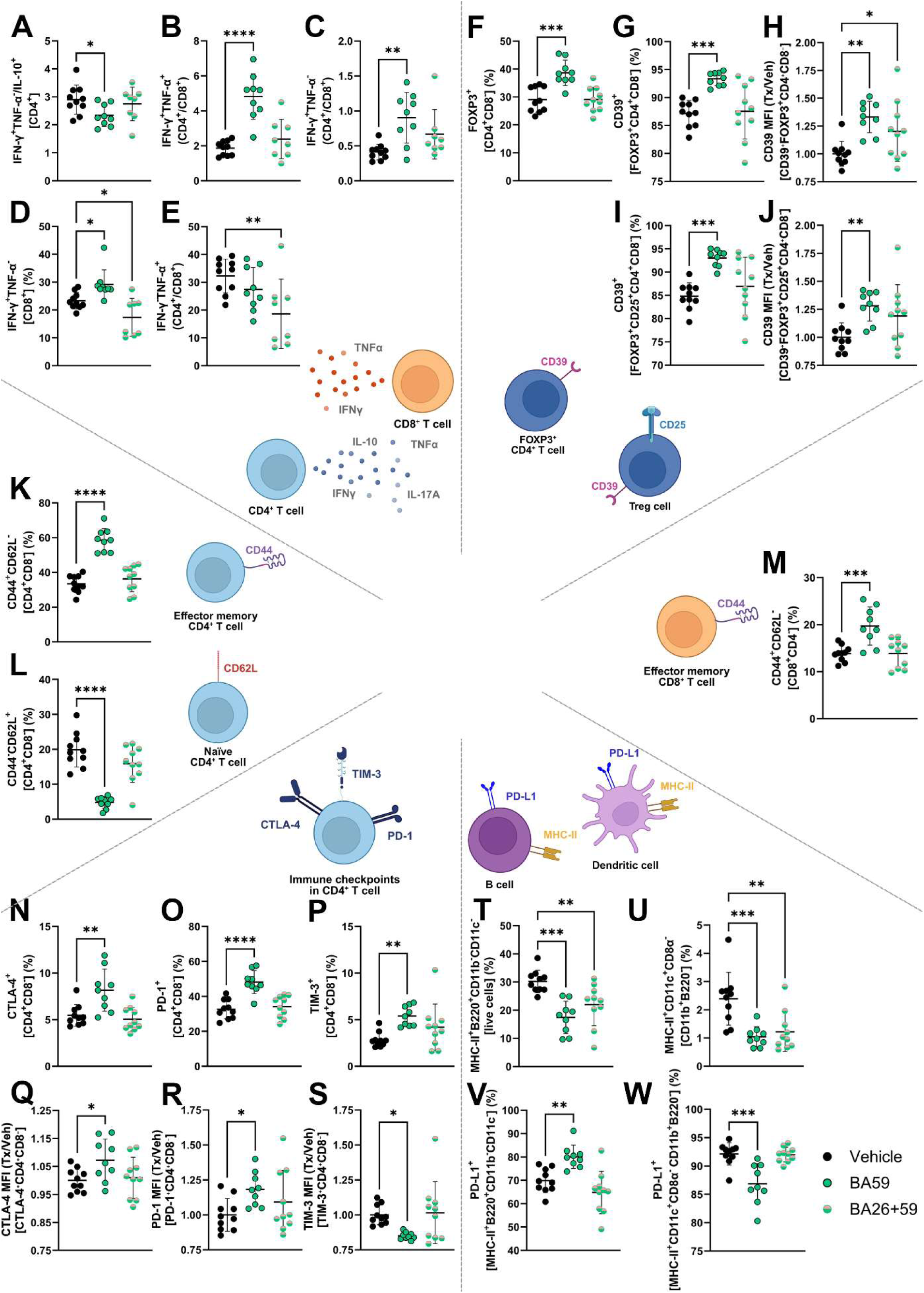
BA59 and combined BA26 and BA59 treatment modulate immune responses across T cell, B cell, and DC compartments in chronic EAE. Multiparameter flow-cytometry analysis was performed using spleen cell suspensions collected at 28 dpi after over 14 days of oral BA treatment. (**A-E**) Cytokine production (IFN-γ, IL-10, IL-17A and TNF-α) by CD4⁺ and CD8⁺ T cells. (**F-J**) Treg cell compartment, showing frequencies of FOXP3⁺ CD4⁺ T cells, CD39⁺ Treg subsets, CD39⁺ Tregs, and CD39 MFI within both CD39⁺ FOXP3⁺ CD4⁺ T cells and CD39⁺ Treg subsets. (**K-M**) T cell memory phenotypes, including effector memory CD4⁺ and CD8⁺ subsets (CD44⁺CD62L⁻) and naïve CD4⁺ frequencies (CD44⁻CD62L⁺). (**N-S**) Expression of key immune checkpoint receptors in CD4⁺ T cells, including CTLA-4, PD-1, and TIM-3, with corresponding MFI analyses. (**T**, **V**) Mature B cell populations (MHC-II⁺B220⁺CD11c⁻CD11b⁻) and PD-L1^+^ frequency within this compartment. (**U**, **W**) Activated mDCs (MHC-II⁺CD11c⁺CD8α⁻CD11b⁺B220⁻) and PD-L1⁺ frequencies within this immune population. Each data point represents an individual mouse, and lines and error bars represent mean ± SD. Vehicle, n=10; BA59, n=9; BA26+59, n=10 (except from cytokine production, n=8). Statistical analysis was performed using one-way ANOVA with Dunnett’s multiple comparison test. * *p value* < 0.05; ** *p value* < 0.01; *** *p value* < 0.001; **** *p value* < 0.0001.

BA59 did not alter the frequency of classical Treg cells (FOXP3⁺CD25⁺) but significantly expanded the broader FOXP3⁺ CD4⁺ population (Figure 4F). Importantly, BA59 enhanced regulatory potential by increasing the proportion of CD39⁺ cells within both FOXP3⁺ and Treg cell subsets and by raising CD39 expression (MFI), indicating higher per-cell expression of this immunosuppressive ectonucleotidase (Figures 4G-J). In contrast, the combined treatment did not modify FOXP3⁺ frequency (Figure 4F) and produced only a modest increase in CD39 expression within FOXP3⁺ cells, suggesting that although it does not expand regulatory populations numerically, it partially enhances their suppressive phenotype through upregulation of CD39 (Figure 4H).

We next assessed how BA treatment influenced CD4⁺ and CD8⁺ T cell differentiation based on CD44 and CD62L expression. BA59 markedly increased the proportion of effector memory T cells (CD44⁺CD62L⁻) in both the CD4⁺ and CD8⁺ compartments, indicating enhanced transition toward antigen-experienced effector states (Figures 4K, M). Consistent with this shift, BA59 produced a pronounced reduction in naïve CD4⁺ T cells (CD44⁻CD62L⁺), reflecting diminished maintenance of the naïve pool (Figure 4L). To further characterize the activation profile of CD4⁺ T cells, we quantified immune-checkpoint expression within CD4⁺ T cells. BA59 increased the frequency of CD4⁺ T cells expressing CTLA-4, PD-1, and TIM-3, indicating broad upregulation of activation-associated inhibitory receptors (Figures 4N-P). Within each checkpoint-positive population, BA59 also enhanced the expression levels (MFI) of CTLA-4 and PD-1 (Figures 4Q, R), while decreasing TIM-3 intensity within TIM-3⁺ CD4⁺ T cells (Figure 4S), suggesting differential regulation of individual checkpoint pathways. In contrast, the combined treatment did not alter the distribution of naïve, effector, or memory T cell subsets (Figures 4K-M), nor did it modify immune checkpoint frequencies or expression levels (Figures 4N-S), indicating that the immunomodulatory effects observed with BA59 alone are not preserved when the two bile acids are co-administered.

Finally, we examined how BA treatment affected APC populations. BA59 alone or in combination with BA26 significantly reduced the frequency of mature B cells defined as MHC-II⁺B220⁺CD11c⁻CD11b⁻ cells (Figure 4T). Within this mature B cell compartment, BA59, but not the combination treatment, increased the proportion of PD-L1⁺ B cells, indicating selective enhancement of regulatory features in BA59-treated mice (Figure 4V). In contrast, the combined treatment did not alter PD-L1 expression frequency, consistent with the loss of immunomodulatory effects observed in other lymphoid subsets. We also evaluated myeloid DCs (mDCs, CD11c⁺CD8α⁻CD11b⁺B220⁻) and found that both BA59 alone and the combination treatment decreased the frequency of activated mDCs (MHC-II⁺) (Figure 4U). However, only BA59 reduced the proportion of PD-L1⁺ activated DCs, suggesting a selective dampening of PD-L1-expressing APCs that was not preserved upon co-administration with BA26 (Figure 4W). Collectively, these results show that BA59 alters both B cell and DC activation states, partially enhancing or suppressing PD-L1 expression depending on the APC subset, whereas the combined treatment abrogates these regulatory shifts.

These findings demonstrate that BA59 exerts a broad reshaping of both adaptive and innate immunity: reducing inflammatory T cell polarization, expanding CD39⁺ regulatory potential in Treg cells, promoting effector memory differentiation, and selectively increasing inhibitory immune checkpoints and PD-L1 pathways. In contrast, co-administration with BA26 consistently abolishes these immunoregulatory effects across both lymphoid and myeloid compartments.

## 4. Discussion

In this study, we systematically evaluated how structural variants of secondary BAs modulate CD4⁺ T cell differentiation and influence autoimmune neuroinflammation. Using a combination of *in vitro* screening and *in vivo* EAE studies, we uncovered distinct immunomodulatory profiles among closely related BA derivatives and identified an androstane-triol derivative (BA59) as a potent modulator of adaptive and innate immune pathways with direct beneficial therapeutic effect in the experimental model of MS.

Our initial single-dose screen demonstrated that both Th17 and Treg cell pathways are highly sensitive to subtle structural differences within the gonane scaffold. Among the 64 tested derivatives, sarsasapogenin (BA26) and an androstane-triol derivative (BA59) emerged as the strongest inhibitors of Th17 cell differentiation, surpassing the positive control dehydroLCA (Figure 2A) [18]. Notably, these two share a hydroxy group at C3 of ring A, a feature previously associated with both Th17 and Treg-modulating capacity, in contrast to the presence of an oxo group at the same position that has been shown to be more selective for Th17 cell polarization (Figure 2C) [18]. In line with that rationale, BA26 showed a dual profile, reducing Th17 cells while increasing Treg cell differentiation, whereas BA59 selectively targeted the Th17 axis (Figure 2A, B). Structural similarities between BA26 and BA49, the latter selectively increasing Treg cell differentiation, highlight how minimal modifications, an additional hydroxy group at C6 in ring B or stereochemistry changes, can invert or enhance specific immunological effects (Figure 2C). Conversely, BA19 reduced both Th17 and Treg cell differentiation while exhibiting cytotoxicity at high doses (50 µM), suggesting that its immunomodulatory profile may partially reflect toxicity linked to its distinct structure features: an oxo substitution at ring C and an acetyl group at ring D (Figure 2C). Since BAs are known for their detergent-like properties that facilitate lipid absorption [16], this effect could partially account for the decrease observed in both immune cell populations in the presence of BA19. These SAR insights expand the structure-function logic previously established for isoalloLCA, dehydroLCA or 3-epideoxycholic acid [18, 19] and emphasize the importance of targeted chemical optimization to bias immune responses toward desired regulatory states.

Dose-response experiments further refined these observations. BA59 exhibited a robust, well-defined sigmoidal response and a lower IC_50_ value (24.04 μM) compared to dehydroLCA (34.24 μM) while maintaining negligible cytotoxicity at 50 μM, the highest tested concentration (Figures 2A, S6). This potency, combined with a clean toxicity profile, supports BA59 as a selective modulator of Th17 cell differentiation positioning it as the most promising derivative for *in vivo* validation. By contrast, most Th17-, Treg-, or dual-acting BAs exhibited sharp declines in cell viability at high concentrations (50 µM), limiting the interpretability of their immunomodulatory effects and constraining their translational potential (Figures S6, S8). Although sarsasapogenin (BA26) exhibited marked cytotoxicity at 50 μM, it was still prioritized for *in vivo* evaluation because of its dual activity and its superior performance relative to both positive controls at 20 μM in both single-dose and dose-response experiments (Figures 1C-D, 2B, S7).

The selective and potent suppression of Th17 cell differentiation by BA59 translated into significant and reproducible therapeutic benefit in chronic EAE. Across independent experiments and dosing paradigms, oral BA59 consistently reduced clinical severity, highlighting its immunomodulatory properties as a disease-modifying candidate (Figures 3B, C). Since profound reductions in BAs have been described in progressive MS patients [20], our results in chronic EAE encourage the translation of this androstane-triol derivative into human testing. Indeed, recent findings from a clinical study in MS patients testing TUDCA, a secondary BA, demonstrated that the compound is safe and biologically active [21], further supporting the feasibility of future clinical trials and the expansion of therapeutic strategies based on gut microbiota-derived metabolites.

On the other hand, despite its *in vitro* activity, BA26 failed to produce clinical improvement in EAE at either 5 mg/kg or 25 mg/kg (Figures S9, S10). This disconnect suggests that its *in vitro* immunomodulatory effects may not be sustained *in vivo*, potentially due to limited bioavailability, physiological instability, or insufficient engagement to relevant immune pathways during neuroinflammation. Thus, these findings illustrate the limitations of relying solely on *in vitro* readouts to predict *in vivo* efficacy.

Comprehensive immunophenotyping identified multiple BA59-driven changes across adaptive and innate compartments (Figure 4). Next, we contextualize the immune shifts that are most strongly connected to EAE immunopathology and most consistent with the clinical effects observed *in vivo*. The treatment with this androstane-triol derivative reduced Th1-skewed CD4⁺ T cell responses and enhanced IL-10-associated regulatory balance. Decreased IFN-γ⁺TNF-α⁻/IL-10⁺ ratios suggest a shift away from pathogenic Th1 polarization (Figure 4A). In contrast to its *in vitro* activity, we did not observe changes in Th17-related ratios with other immunoregulatory (IL-10⁺/IL-17A⁺) or pro-inflammatory T cell subsets (IFN-γ⁺TNF-α⁻/IL-17A⁺ and IFN-γ⁺TNF-α⁺/IL-17A⁺) in the periphery (Figure S12). This may indicate that, although the androstane-triol derivative reshapes other immune compartments, its Th17-modulatory effects do not extend beyond the gut-associated immune environment. Notably, other secondary BAs with known Th17-modulatory properties often exert their effects in the gut or draining lymph nodes, with limited evidence for systemic modulation [18, 22]. Because we did not analyze gut tissues, less relevant to our disease model, we cannot determine whether local Th17 cell modulation occurred. However, our data clearly show that peripheral and systemic Th17 cell responses were not affected.

BA59 remodeled the effector T cell compartment toward a less cytotoxic profile. In treated mice, the CD4⁺/CD8⁺ effector balance shifted toward CD4⁺ T cells, while CD8⁺ T cells were skewed toward IFN-γ single producers rather than IFN-γ/TNF-α double producers (Figures 4B-D). Although total *bona fide* Treg cell frequencies (FOXP3^+^ CD25^+^CD4^+^) remained stable, BA59 markedly increased broader FOXP3^+^CD4^+^ Treg cell population as well as CD39⁺ populations within the Treg and FOXP3^+^ cell compartments. CD39 is required for ATP hydrolysis and contributes to adenosine-mediated suppression, and increased CD39 expression has been associated with enhanced regulatory function in autoimmune settings [26–28]. Additionally, this androstane-triol derivative also upregulated CD39 expression per cell in both regulatory CD4^+^ T cell populations, indicating a qualitative enhancement of regulatory function.

Together with the expansion of CD44⁺CD62L⁻ effector-memory CD4⁺ and CD8⁺ T cell subsets, these data suggest that BA59 favors the formation of an antigen-experienced T cell pool with attenuated cytotoxicity, which is compatible with durable control of neuroinflammation rather than ongoing destructive effector activity.

BA59 treatment also increased the CD4^+^ T cell populations expressing the inhibitory receptors CTLA-4, PD-1, and TIM-3 on CD4^+^ T cells and, to a certain extent, also an increment on their expression levels per cell, indicating enhanced engagement of negative regulatory pathways that are typically diminished in MS (Figures 4N-S) [29, 30]. In EAE, loss or blockade of any of these checkpoints consistently increases frequency of pro-inflammatory-expressing T cells, promotes CNS infiltration, and accelerates disease, underscoring their central role in restraining autoimmune neuroinflammation [31–34]. The checkpoint upregulation observed with BA59 parallels the reduction in IFN-γ⁺TNF-α⁻/IL-10 ratios, suggesting a coordinated shift toward less inflammatory effector programs and a microenvironment more permissive to IL-10-mediated regulation. These immunological adjustments collectively provide a mechanistic framework linking BA59-induced checkpoint restoration with the attenuation of Th1-driven pathology and the clinical improvement observed in treated EAE mice At the APC level, BA59 reduced mature B cell frequencies while increasing PD-L1⁺ regulatory B cells and decreased activated mDCs, while altering PD-L1 expression patterns (Figures 4T-W). These broad and coordinated shifts suggest that BA59 rebalances antigen presentation and costimulatory signals to favor a more tolerogenic immune environment during chronic neuroinflammation.

A key observation is that the combined treatment with both BA26 and BA59 produced clinical improvement comparable to BA59 alone (Figure 3C) yet failed to reproduce most of the observed immunological signatures of the single treatment (Figure 4). The combination barely modified cytokine profiles, Treg cell subsets, and APC phenotypes; and had no effect on immune checkpoint expression and effector-memory differentiation. These findings may be indicating functional antagonism between BA26 and BA59 despite structural similarity. The most plausible explanations include competition for shared BA receptors (e.g., FXR, GPBAR1) [35], interference with oral bioavailability and tissue distribution, or induction of opposing transcriptional programs. That clinical improvement persists despite immunological discrepancies highlights the complexity of BA-mediated signaling *in vivo* and suggests that clinical readouts may not always reflect underlying mechanistic pathways.

Together, our data reveal that BA59 acts as a potent and selective immunomodulatory BA derivative capable of reprogramming key immune circuits in chronic EAE. Its effects span Th1/Treg cell balance, regulatory CD39⁺ subsets, effector-memory differentiation, immune checkpoint regulation, and APC activation states. In contrast, BA26 lacks *in vivo* efficacy and disrupts BA59-driven immunoregulation when co-administered.

Overall, these findings underscore three major principles for BA-based immunotherapy development: i) minor structural modifications produce major functional consequences, necessitating precise SAR-guided optimization; ii) *in vitro* potency does not guarantee *in vivo* activity, highlighting the need to integrate receptor-engagement and pharmacokinetic data early on in the development process, and iii) combinatorial use of structurally related BAs may produce antagonistic interactions, underscoring the importance of understanding shared pathways and potential crosstalk between compounds.

Future work should define the molecular targets, receptor affinities and metabolic processing of both BA candidates to elucidate the mechanistic basis of their divergent effects. Overall, BA59, an androstane-triol derivative, emerges as a promising compound and lead scaffold for designing next-generation BA derivatives tailored to rebalance pathogenic immune circuits in CNS autoimmunity.

## 5. Conclusions

Our work demonstrates that selective chemical modifications of the BA scaffold gonane can profoundly influence not only T cell compartment but also APC populations and disease outcomes in EAE. BA59 emerges as a potent anti-inflammatory candidate capable of reshaping immune networks across both adaptive and innate compartments. In contrast, BA26 exerts strong *in vitro* activity but lacks *in vivo* efficacy and disrupts the immunoregulatory effects of BA59 when co-administered. These insights provide a foundation for rational design of BA-based therapeutics and underscore the value of integrating SAR, immunophenotyping, and *in vivo* functional studies to identify derivatives with optimal therapeutic profiles.

## Acknowledgements

We used instruments supported by the Flow Cytometry Core Facility (Weill Cornell Medicine) for our *in vitro* experiments. We thank Mireia Castillo and Alba Hernández-Badosa for their valuable technical support in the *in vivo* and *ex vivo* studies performed in the EAE model.

## Authors’ contributions

MTG and LCB conceived and designed the study. LCB performed the *in vitro* T-cell differentiation experiments. The EAE *in vivo* experiments and *ex vivo* immune response studies were conducted by IBB, HE, and CE. IBB performed CNS stainings and acquired data for the histopathological studies. LCB analyzed all datasets and prepared the figures. LCB wrote the first draft of the manuscript, and IBB, HE, CE, and MTG contributed to manuscript revisions. LCB, CE, and MTG supervised the work.

MTG and CE secured the funding that supported this work. All authors read and approved the final manuscript.

## Funding

Support was provided by the Agència de Gestió d’Ajuts Universitaris i de Recerca (AGAUR; Generalitat de Catalunya) through the Consolidated Research Groups program (2021SGR00782).

## Availability of data and materials

Most of the datasets generated and analyzed during the current study are included in this article and its supplementary information files. Additional raw data not shown in the manuscript, such as full flow cytometry FCS files, complete histopathology image sets, and other primary experimental outputs, are available from the corresponding author upon reasonable request.

## Declarations

### Ethics approval and consent to participate

All animal procedures were conducted in accordance with the ARRIVE guidelines and the NIH *Guide for the Care and Use of Laboratory Animals*. For in vitro experiments, male and female C57BL/6J mice (5-6 weeks old; Jackson Laboratory, Stock #000664) were housed under standard light- and climate-controlled conditions with ad libitum access to food and water. All procedures were approved by the Institutional Animal Care and Use Committee (IACUC) at Weill Cornell Medicine (protocol number: 2022-0040).

For *in vivo* EAE studies, female C57BL/6JOlaHsd mice (8 weeks old; Envigo, Venray, The Netherlands) were housed under standard conditions with ad libitum access to chow and water. Experiments were conducted in compliance with European Union regulations (Directive 2010/63/EU) and Spanish legislation (Real Decreto 53/2013; Generalitat de Catalunya Decret 214/97). All procedures were approved by the Ethics Committee on Animal Experimentation of the Vall d’Hebron Research Institute (protocol number: CEEA 46/19: CEA-OH/9459R1/2). All data presented are in accordance with the guidelines suggested for EAE publications [23] and with the ARRIVE guidelines for animal research.

No human participants or human-derived materials were used in this study.

## Consent for publication

Not applicable.

## Competing interests

The authors declare that they have no competing interests.

## List of abbreviations

AUC: area under the curve
APC: antigen-presenting cell
ARRIVE: Animal Research: Reporting of In Vivo Experiments
BA: bile acid
CNS: central nervous system
DAPI: 4’,6-diamidino-2-phenylindole
DCA: deoxycholic acid
DC: dendritic cell
Dpi: day post-immunization
EAE: experimental autoimmune encephalomyelitis
FMO: fluorescence minus one
FXR: farnesoid X receptor
H&E: haematoxylin and eosin
HLA: human leukocyte antigen
IFA: incomplete Freund’s adjuvant
IACUC: Institutional Animal Care and Use Committee
LCA: lithocholic acid
mAb: monoclonal antibody
mDC: myeloid dendritic cell
MFI: median fluorescence intensity
MS: multiple sclerosis
PMA: phorbol 12-myristate 13-acetate
SAR: structure-activity relationship
Th: T helper
Treg: regulatory T
TUDCA: tauroursodeoxycholic acid
4PL: four-parameter logistic

## Notes

### Competing Interest Statement

The authors have declared no competing interest.

