## Supporting Information for "Bile Acid Scaffold Engineering Reveals an Androstane-Triol Derivative as a Potent Immunomodulator with Therapeutic Efficacy in EAE"

### 1. Supplementary Tables

**Table S1. Compounds included in the bile acid screening library.** List of 66 compounds used in the bile acid screening, including compound IDs as referenced throughout the manuscript, compound name, catalog numbers, and vendor names.

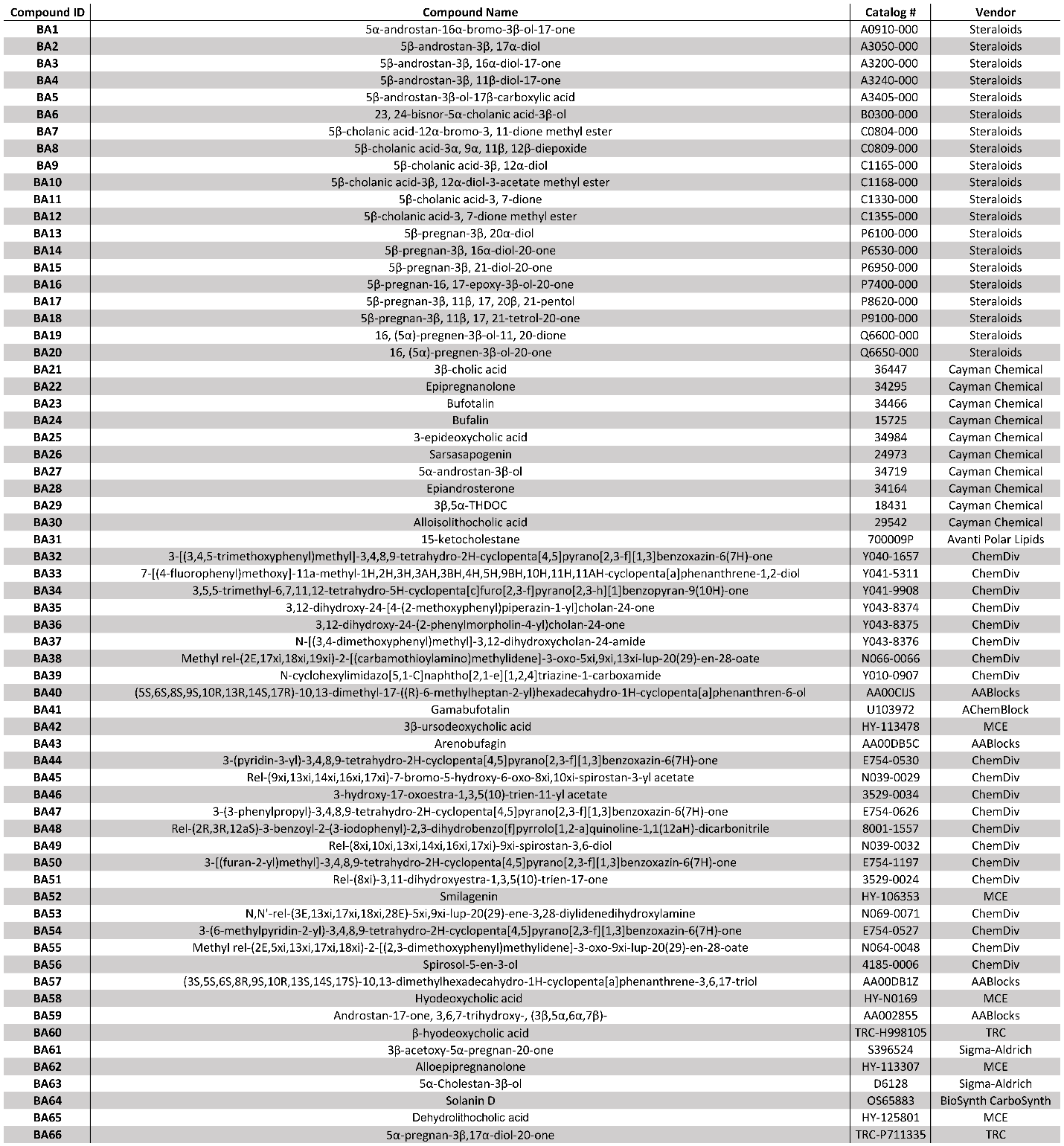

### 2. Supplementary Figures

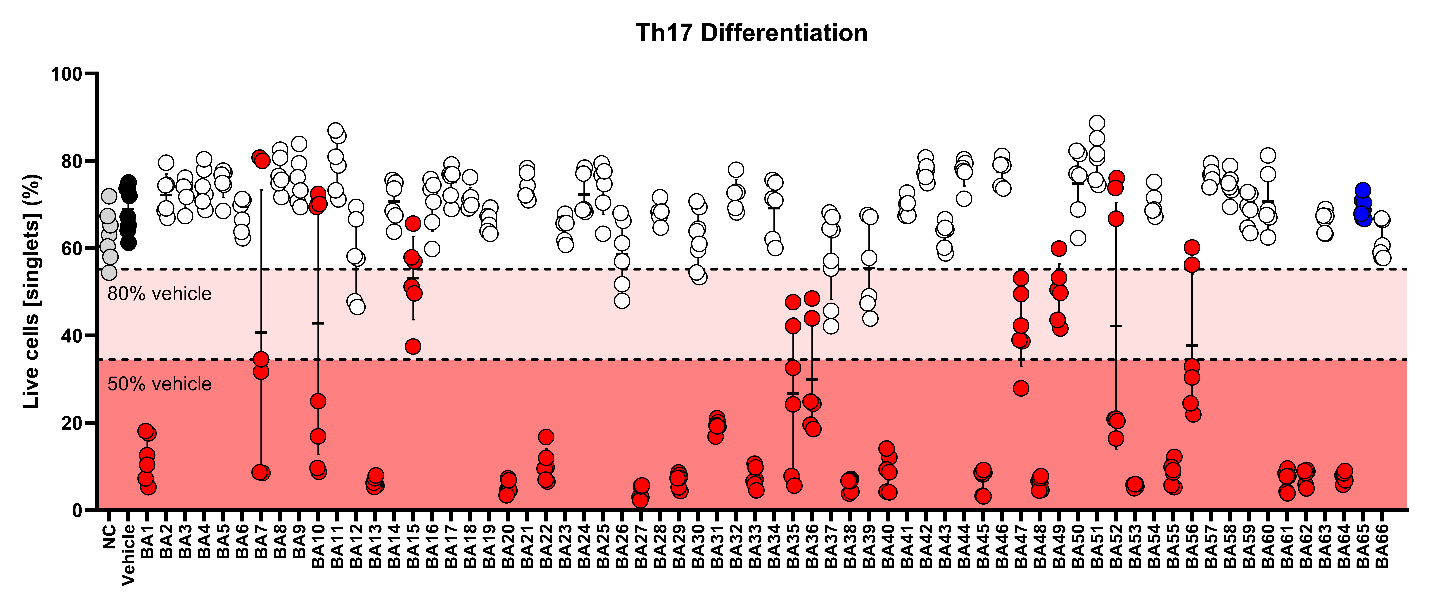

**Figure S1. Effect of screened compounds on cell viability under Th17-differentiating conditions.** The positive control, BA65 (dehydrolithocholic acid), is shown in dark blue. The negative control (NC, Th17-differentiating conditions without DMSO) is shown in grey and included to assess the effect of DMSO alone. The vehicle control (Th17-differentiating conditions with DMSO at concentration 0.4%) is shown in black and serves as the reference condition for evaluating compound effects on cell viability. Samples shown in red indicate a statistically significant reduction in viability compared to the vehicle control (*p value* < 0.05). The dark red shaded area marks samples with viability reduced by more than 50%, while the light red shaded area indicates a reduction between 20% and 50%. No significant difference was observed between the negative and vehicle controls. The experimental data represents four independent experiments with six to eight replicates per experimental condition. Individual values for each replicate are displayed, along with the mean and standard deviation across all samples. Statistical analyses were performed using one-way ANOVA with Dunnett's multiple comparison test.

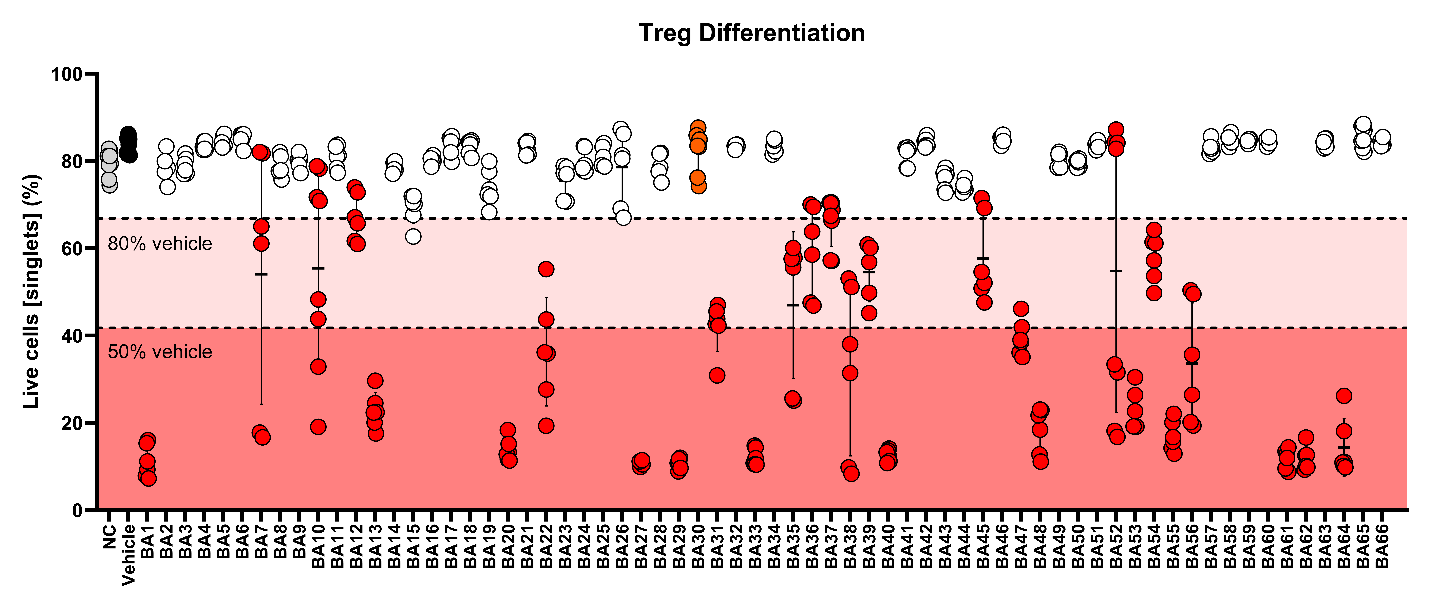

**Figure S2. Effect of screened compounds on cell viability under Treg-differentiating conditions.** The positive control, BA30 (alloisolithocholic acid), is shown in dark orange. The negative control (NC, Treg-differentiating conditions without DMSO) is shown in grey and included to assess the effect of DMSO alone. The vehicle control (Treg-differentiating conditions with DMSO at concentration 0.4%) is shown in black and serves as the reference condition for evaluating compound effects on cell viability. Samples shown in red indicate a statistically significant reduction in viability compared to the vehicle control (*p value* < 0.05). The dark red shaded area marks samples with viability reduced by more than 50%, while the light red shaded area indicates a reduction between 20% and 50%. No significant difference was observed between the negative and vehicle controls. The experimental data represents four independent experiments with six to eight replicates per experimental condition. Individual values for each replicate are displayed, along with the mean and standard deviation across all samples. Statistical analyses were performed using one-way ANOVA with Dunnett's multiple comparison test.

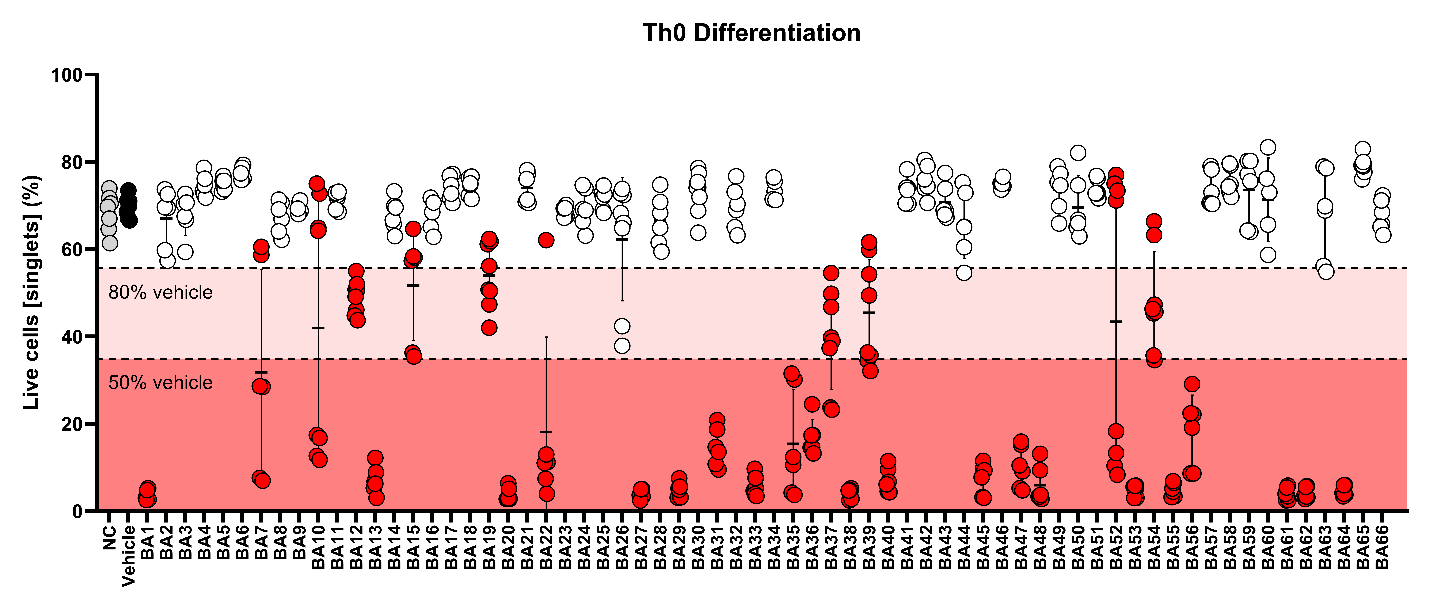

**Figure S3. Effect of screened compounds on cell viability under Th0-differentiating conditions.** The negative control (NC, Th0-differentiating conditions without DMSO) is shown in grey and included to assess the effect of DMSO alone. The vehicle control (Th0-differentiating conditions with DMSO at concentration 0.4%) is shown in black and serves as the reference condition for evaluating compound effects on cell viability. Samples shown in red indicate a statistically significant reduction in viability compared to the vehicle control (*p value* < 0.05). The dark red shaded area marks samples with viability reduced by more than 50%, while the light red shaded area indicates a reduction between 20% and 50%. No significant difference was observed between the negative and vehicle controls. The experimental data represents four independent experiments with six to eight replicates per experimental condition. Individual values for each replicate are displayed, along with the mean and standard deviation across all samples. Statistical analyses were performed using one-way ANOVA with Dunnett's multiple comparison test.

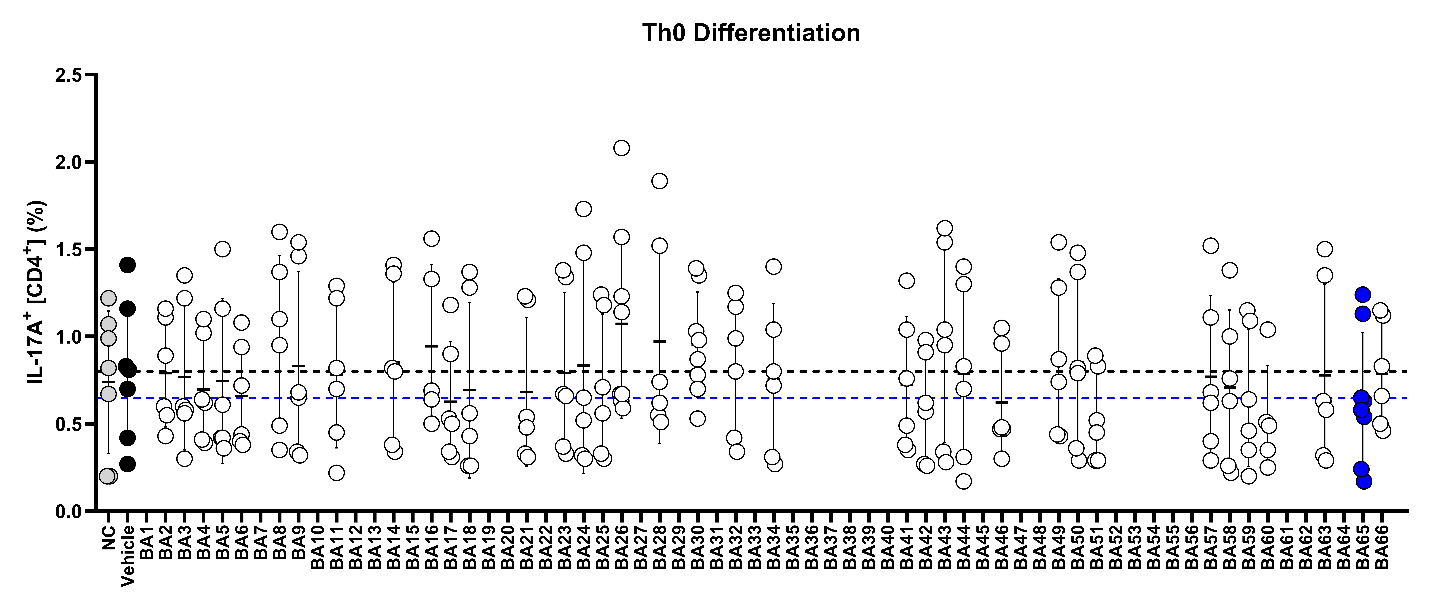

**Figure S4. Effect of screened compounds on IL-17A-expressing cells under Th0-differentiating conditions.** The positive control, BA65 (dehydrolithocholic acid), is represented in dark blue. The negative control (NC, Th0-differentiating conditions without DMSO) is shown in grey and was included to assess the baseline effect without the presence of DMSO. The vehicle control (Th0-differentiating conditions with DMSO at concentration 0.4%) is shown in black and serves as the reference condition for evaluating compound effects on IL-17A expression. No screened compound induced a statistically significant change in IL-17A expression compared to the vehicle control (*p value* < 0.05). Additionally, no significant difference was observed between the negative and vehicle controls. The black dashed line indicates the mean IL-17A expression of the vehicle control, while the dark blue dashed line indicates the mean expression of the positive control. The experimental data represents four independent experiments with six to eight replicates per experimental condition. Individual values for each replicate are displayed, along with the mean and standard deviation across all samples. Statistical analyses were performed using one-way ANOVA with Dunnett's multiple comparison test.

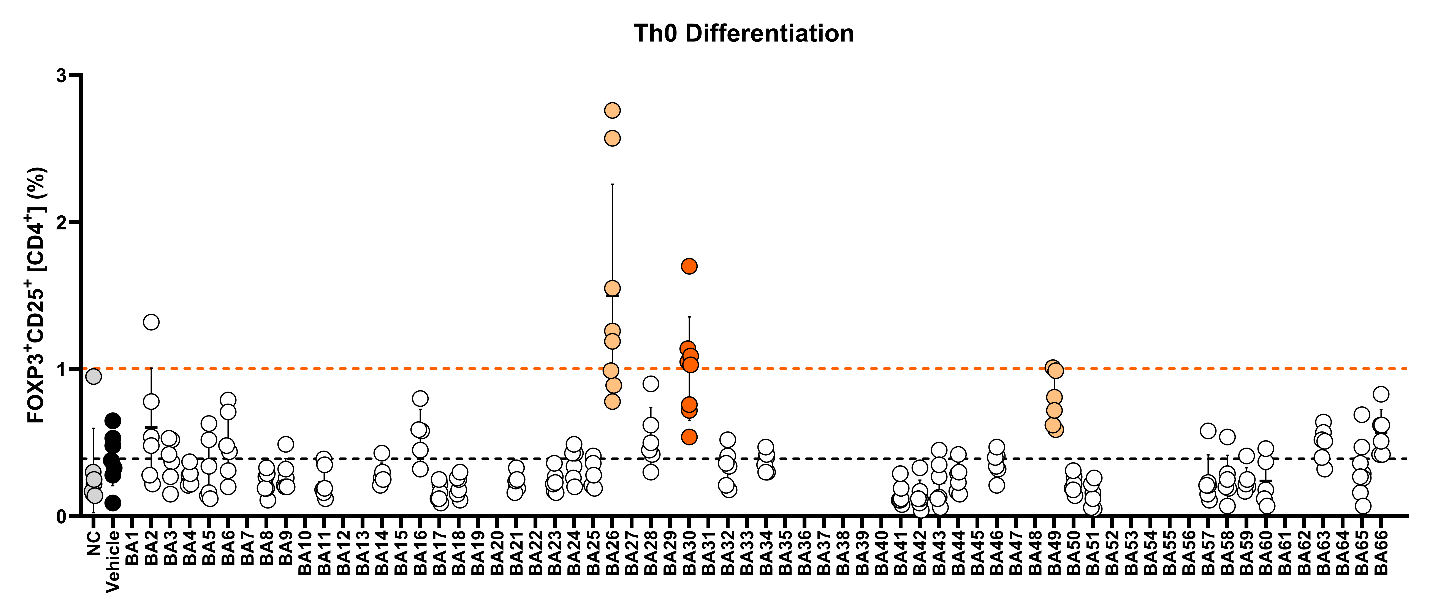

**Figure S5. Effect of screened compounds on Treg cells under Th0-differentiating conditions.** The positive control, BA30 (alloisolithocholic acid), is represented in dark orange. The negative control (NC, Th0-differentiating conditions without DMSO) is shown in grey and was included to assess the baseline effect without the presence of DMSO. The vehicle control (Th0-differentiating conditions with DMSO at concentration 0.4%) is shown in black and serves as the reference condition for evaluating compound effects on Treg cells. Both the positive control as well as those compounds in light orange color showed a statistically significant change in Treg cell frequencies compared to the vehicle control (*p value* < 0.05). No significant difference was observed between the negative and vehicle controls. The black dashed line indicates the mean Treg cell frequency of the vehicle control, while the dark orange dashed line indicates the mean frequency of the positive control. The experimental data represents four independent experiments with six to eight replicates per experimental condition. Individual values for each replicate are displayed, along with the mean and standard deviation across all samples. Statistical analyses were performed using one-way ANOVA with Dunnett's multiple comparison test.

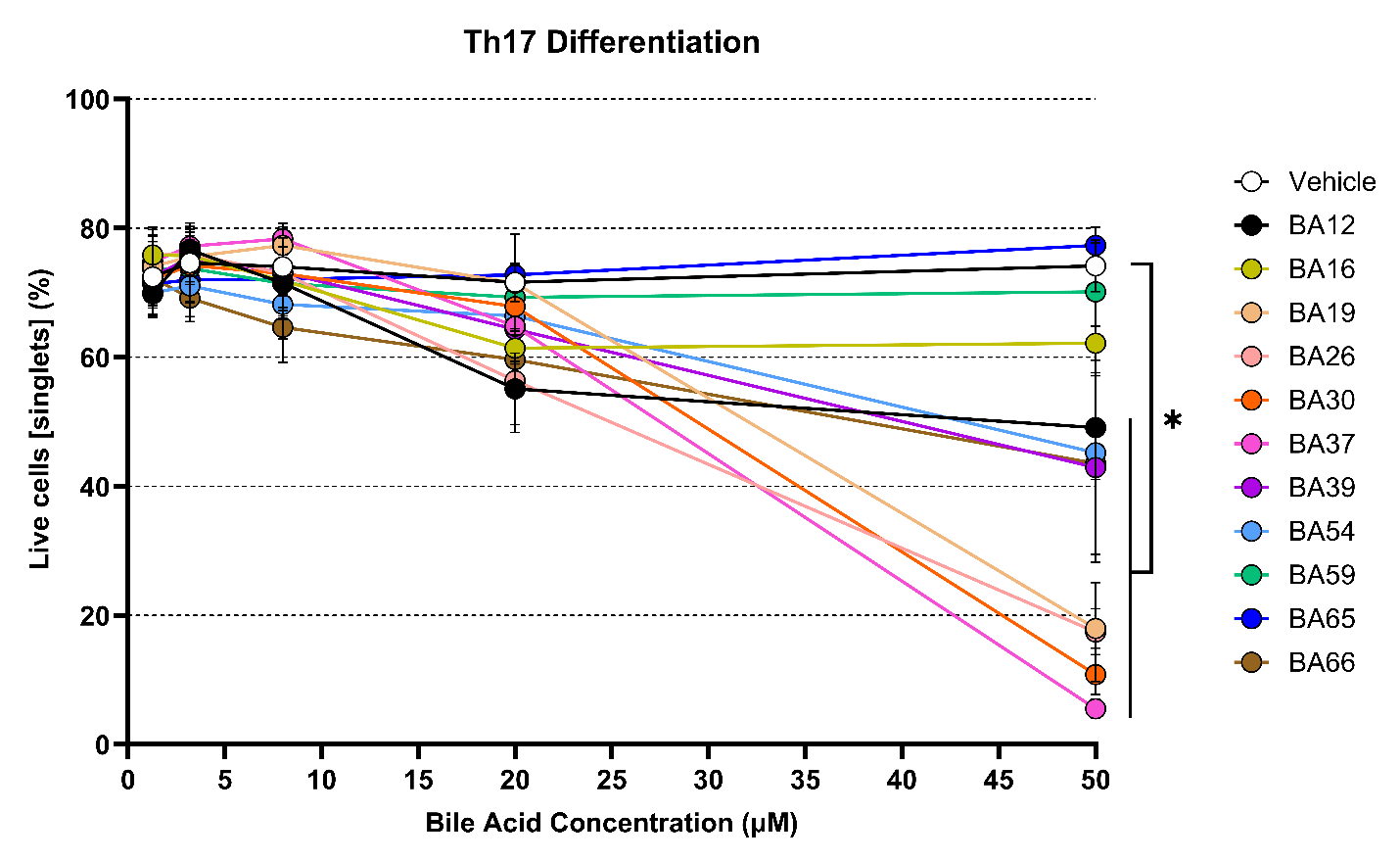

**Figure S6. Dose-response analysis of screened compounds on cell viability under Th17-differentiating conditions.** Each compound was tested at five concentrations: 50 µM, 20 µM (same as in the single-dose screening), 8 µM, 3.2 µM, and 1.28 µM. The vehicle control (Th17-differentiating conditions with DMSO at a constant concentration of 0.4%) serves as the reference for evaluating compound effects. The positive control, BA65 (dehydrolithocholic acid), is shown in dark blue and did not affect cell viability at any tested concentration compared to the vehicle. Compounds BA12, BA19, BA26, BA30, BA37, BA39, BA54, and BA66 significantly reduced cell viability at 50 µM (*p value* < 0.05). No statistically significant differences were observed at lower concentrations. Data represent three independent experiments, each with one technical replicate. Mean values and standard deviations for each concentration are shown. Statistical analyses were performed using one-way ANOVA with Dunnett's multiple comparison test. * *p value* < 0.05.

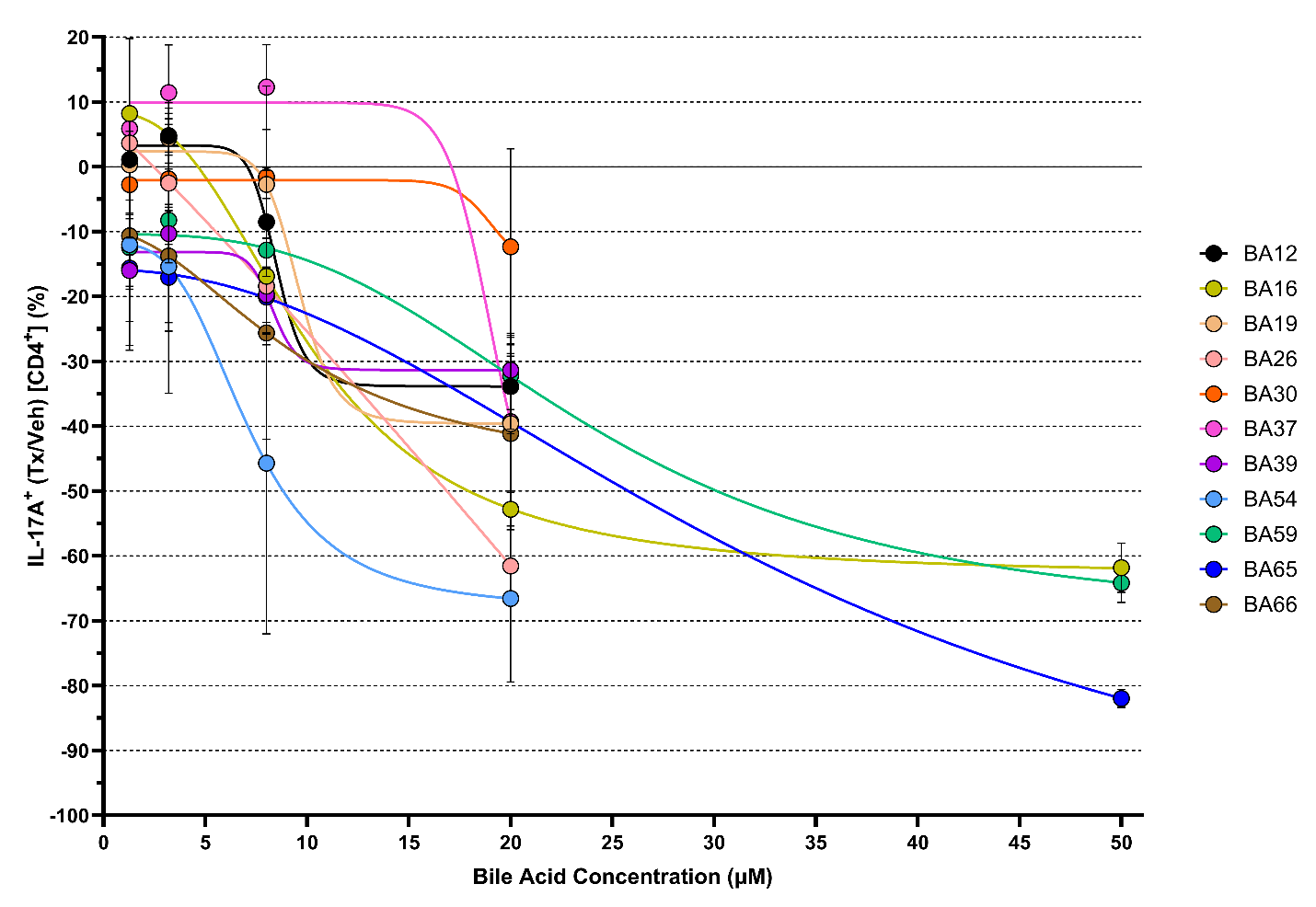

**Figure S7. Selected BA derivatives exhibit dose-dependent effects on Th17 populations.** Naïve CD4^+^ T cells were cultured under Th17-polarizing conditions and treated with increasing concentrations: 1.28 µM, 3.2 µM, 8 µM, 20 µM (same as in the single-dose screening), and 50 µM; of previously identified active BAs (BA12, BA16, BA19, BA26, BA30, BA37, BA39, BA54, BA59, and BA66) or positive control (BA65, dehydroLCA). All tested compounds, as well as the positive control (BA65), reduced Th17 population frequency in a dose-dependent manner within the non-toxic range for each candidate compound. Data represent three independent experiments, each with one technical replicate. Mean values and standard deviations for each concentration are shown. IC_50_ values were calculated by fitting dose-response data to a sigmoidal four-parameter logistic (4PL) model, using BA concentration as the X variable. IC_50_ values were derived from 5-point dose-response curves; when compounds exhibited reduced viability at the highest concentration, only the four non-toxic concentrations were included in the fit.

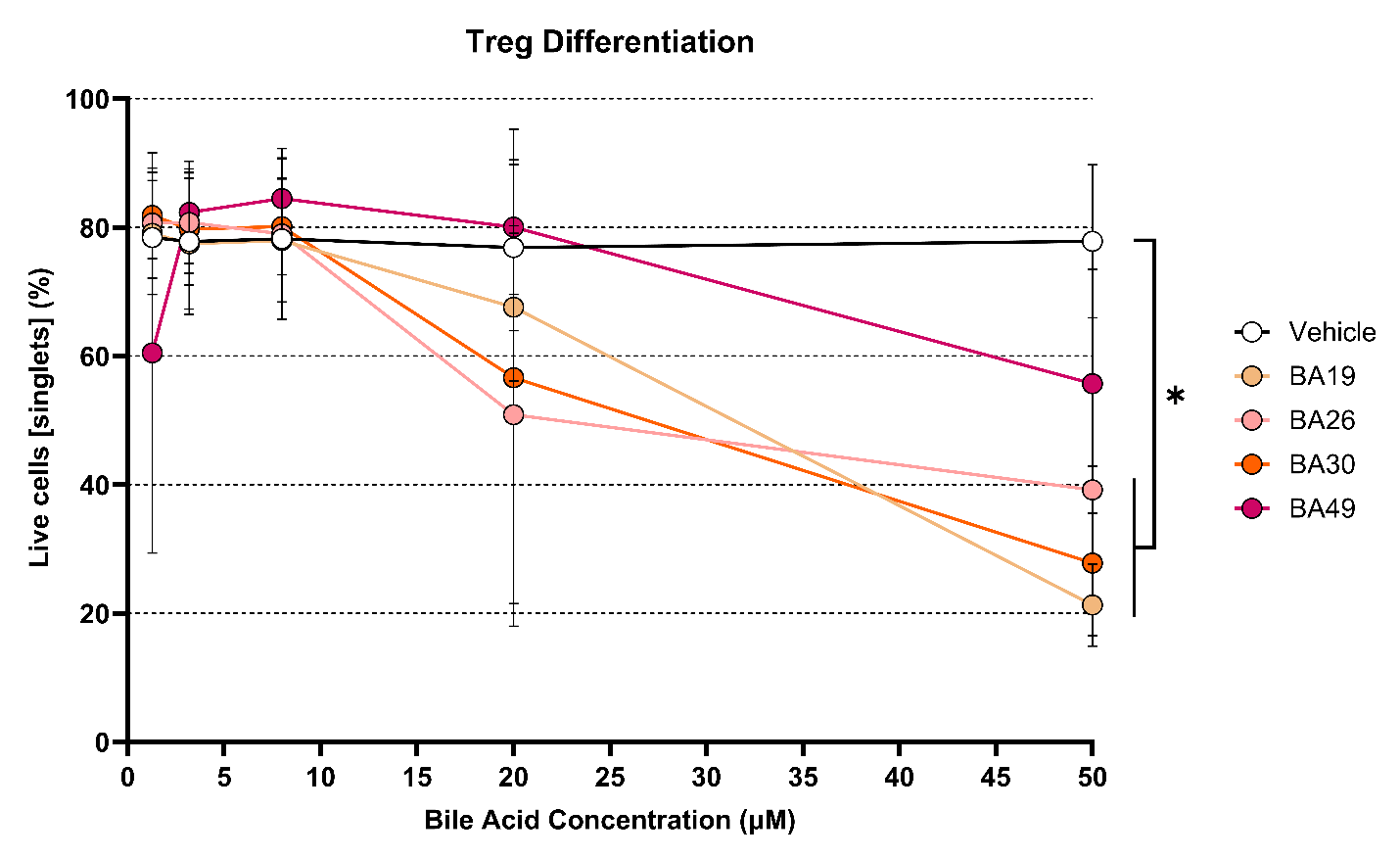

**Figure S8. Dose-response analysis of screened compounds on cell viability under Treg-differentiating conditions.** Each compound was tested at five concentrations: 50 µM, 20 µM (same as in the single-dose screening), 8 µM, 3.2 µM, and 1.28 µM. The vehicle control (Treg-differentiating conditions with DMSO at a constant concentration of 0.4%) serves as the reference for evaluating compound effects. The positive control, BA30 (isoalloLCA), is shown in dark orange and reduced cell viability at 50 µM compared to the vehicle (*p value* < 0.05). Moreover, compounds BA19 and BA26 significantly reduced cell viability at 50 µM (*p value* < 0.05). No statistically significant differences were observed at lower concentrations. Data represent three independent experiments, each with one technical replicate. Mean values and standard deviations for each concentration are shown. Statistical analyses were performed using one-way ANOVA with Dunnett's multiple comparison test. * *p value* < 0.05.

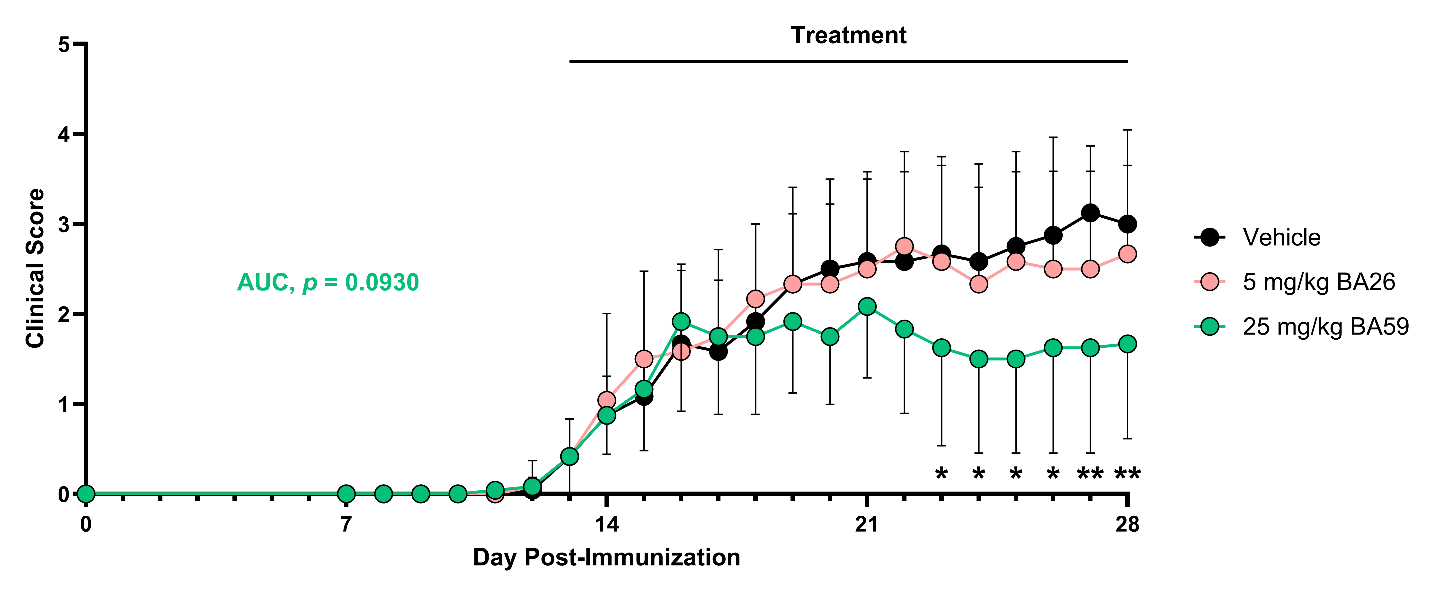

**Figure S9. BA59 reduced the clinical severity of EAE in a therapeutic approach.** Daily oral administration of BA59 (25 mg/kg), initiated when mice reached a clinical score equal to or higher than 1, resulted in an improvement in EAE clinical outcomes. In contrast, daily oral administration of BA26 (5 mg/kg) did not confer clinical benefit. The graph represents the mean clinical score throughout the EAE clinical course of a single experiment (Vehicle, n=12; BA26, n=12; and BA59, n=12). The data are presented as the means and standard deviations. Statistical analyses were performed using one-way ANOVA with Dunnett’s multiple comparison test (Vehicle as the control group) for both analyses: (i) clinical scores compared across groups at each day post-immunization, and (ii) area-under-the-curve (AUC) values calculated for each mouse and compared between groups. * *p value* < 0.05; ** *p value* < 0.01.

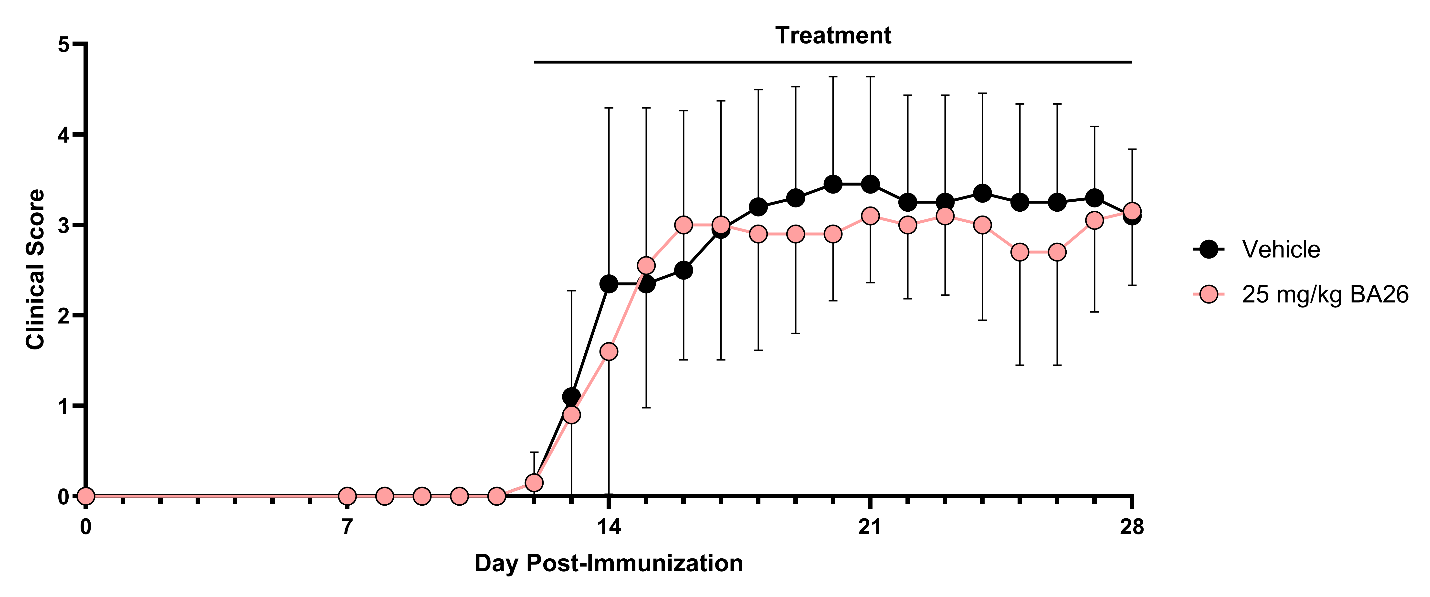

**Figure S10. BA26 did not reduce the clinical severity of EAE in a therapeutic approach.** Daily oral administration of BA26 (25 mg/kg), initiated when mice reached a clinical score equal to or higher than 1, did not exert a clinical improvement in EAE mice. The graph represents the mean clinical score throughout the EAE clinical course of a single experiment (Vehicle, n=10; and BA26, n=10). The data are presented as the means and standard deviations.

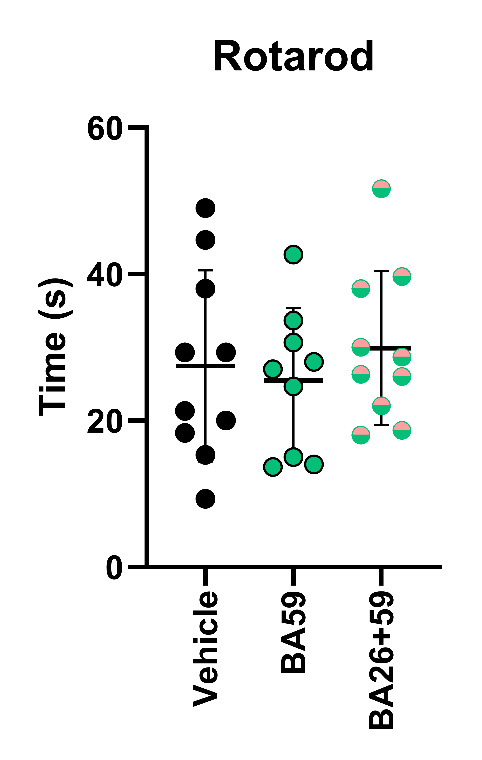

**Figure S11. Treatment with BA59 alone or in combination with BA26 did not improve motor function in EAE mice.** Daily oral administration of BA59 (25 mg/kg) or BA26+BA59 (25 mg/kg each), initiated when mice reached a clinical score equal to or higher than 1, did not improve motor function at the study endpoint (28 dpi), as assessed by the Rotarod test. The graph shows the mean latency to fall (three technical replicates per animal) from a single experiment (Vehicle, n=10; and BA59, n=9; BA26+59, n=10). Data are also presented as mean ± SD for each experimental group.

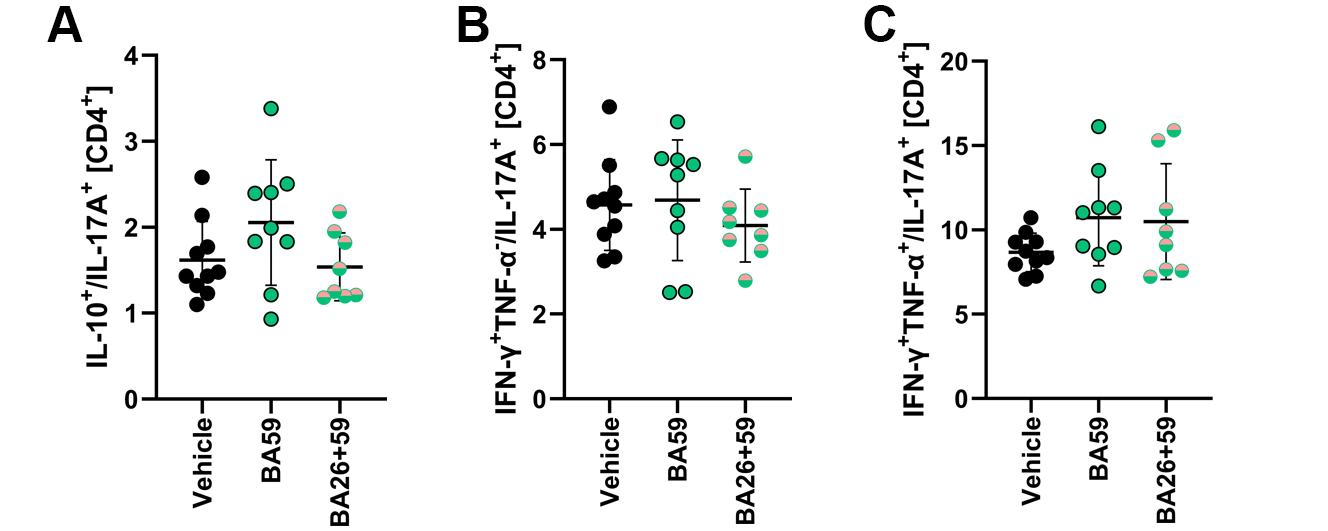

**Figure S12. Treatment with BA59 alone or in combination with BA26 did not alter peripheral IL-17A⁺ T-cell ratios.** Daily oral administration of BA59 (25 mg/kg) or BA26+BA59 (25 mg/kg each), initiated when mice reached a clinical score equal to or higher than 1, did not change the ratios between Th17 populations (IL-17A^+^) and other (**A**) immunoregulatory (IL-10^+^) and (**B**, **C**) pro-inflammatory (IFN-γ^+^TNF-α^-^, IFN-γ^+^TNF-α^+^) populations at the study endpoint (28 dpi). The graphs show individual ratios per animal from a single experiment (Vehicle, n=10; and BA59, n=9; BA26+59, n=10). Data are also presented as mean ± SD for each experimental group.

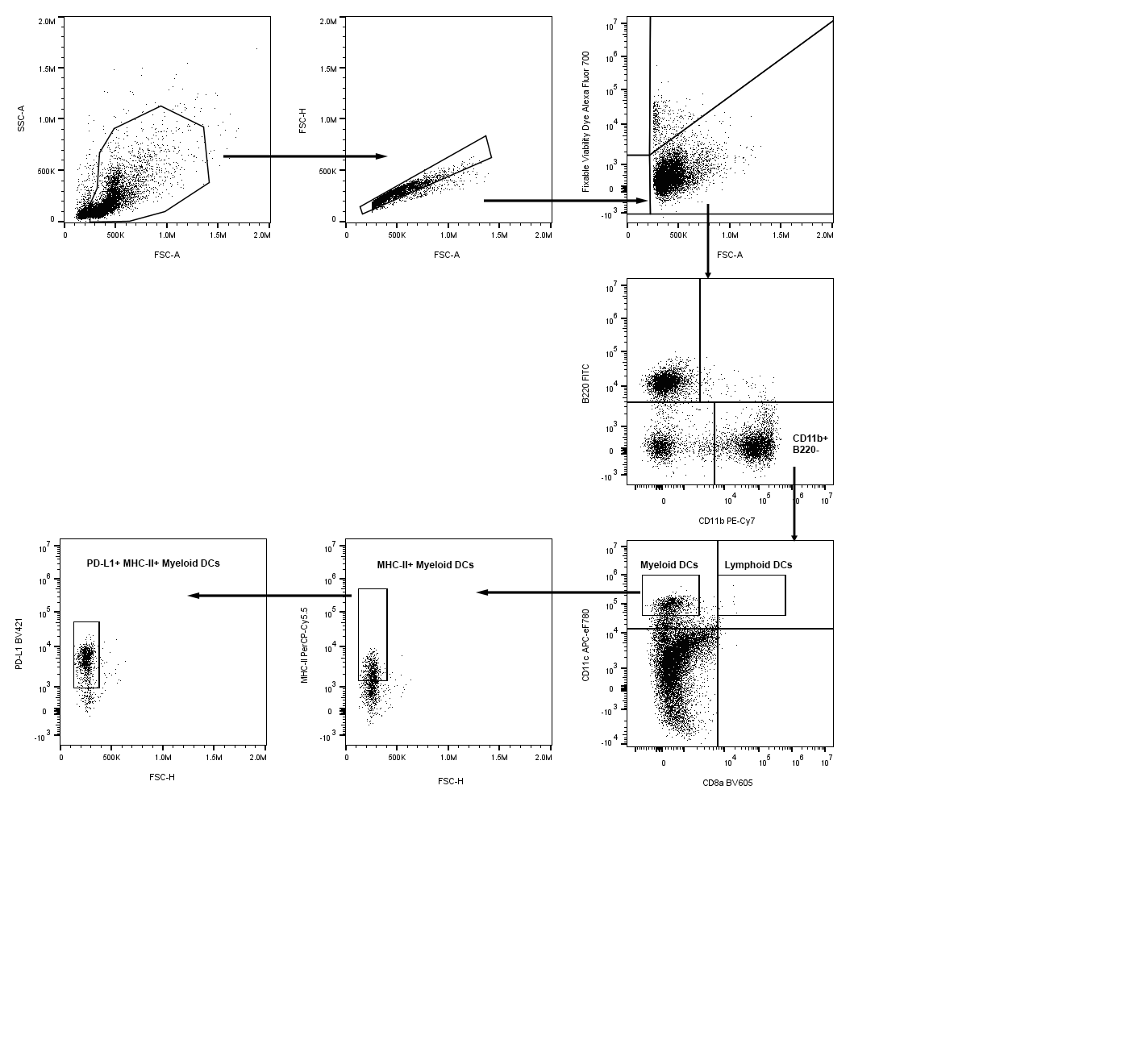

**Figure S13. Flow cytometry gating strategy for DC analysis.** Viable cells were first identified by exclusion of staining with a fixable viability dye. Subsequent gating steps were applied to define DC subsets and assess the expression of immune receptors MHC-II and PD-L1 within the myeloid DC population.

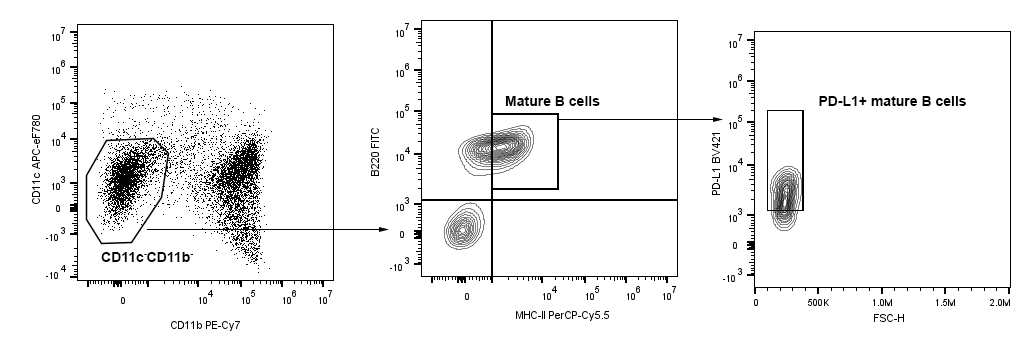

**Figure S14. Flow cytometry gating strategy for B cell cell analysis.** Viable cells were first identified by exclusion of staining with a fixable viability dye (as described in *Figure S13*). Subsequent gating steps were applied to define B cell (B220^+^CD11c^-^CD11b^-^) and assess the expression of MHC-II and PD-L1 within this population.

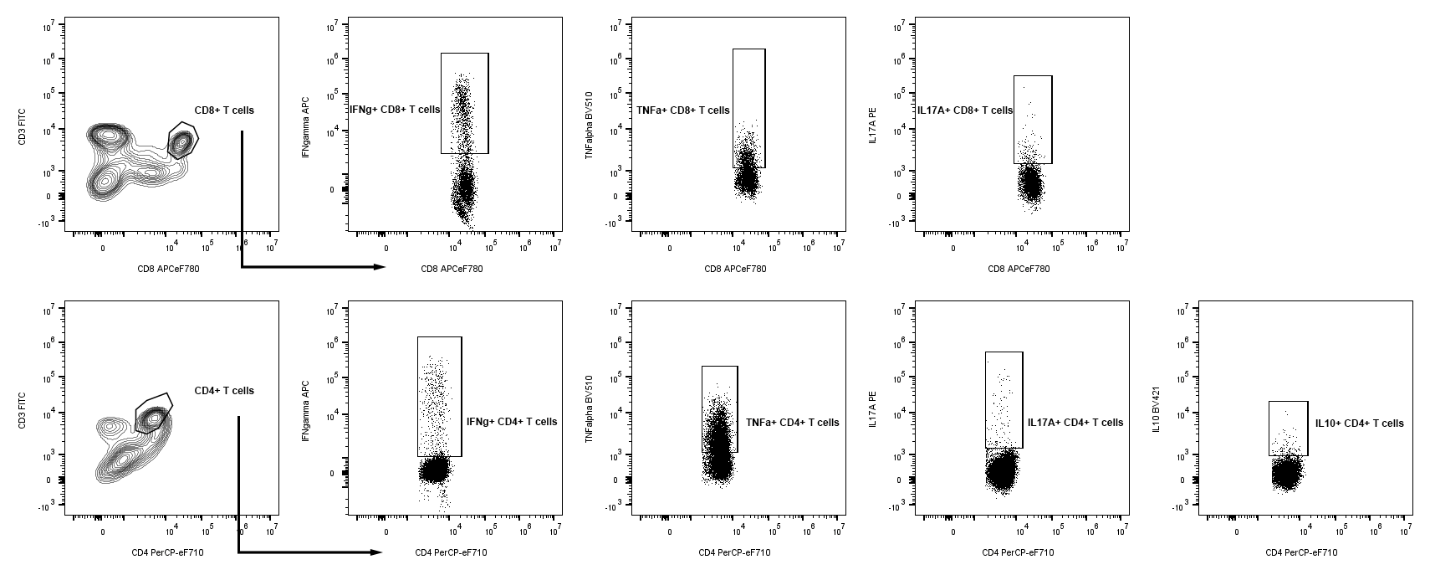

**Figure S15. Flow cytometry gating strategy for cytokine expression in T cells.** Viable cells were first identified by exclusion of staining with a fixable viability dye (as described in *Figure S13*). Subsequent gating steps were applied to analyze the expression of cytokines: including interferon gamma (IFN-γ), tumor necrosis factor alpha (TNF-α), interleukin 17A (IL-17A), and interleukin 10 (IL-10); in CD4⁺ and/or CD8⁺ T cell populations.

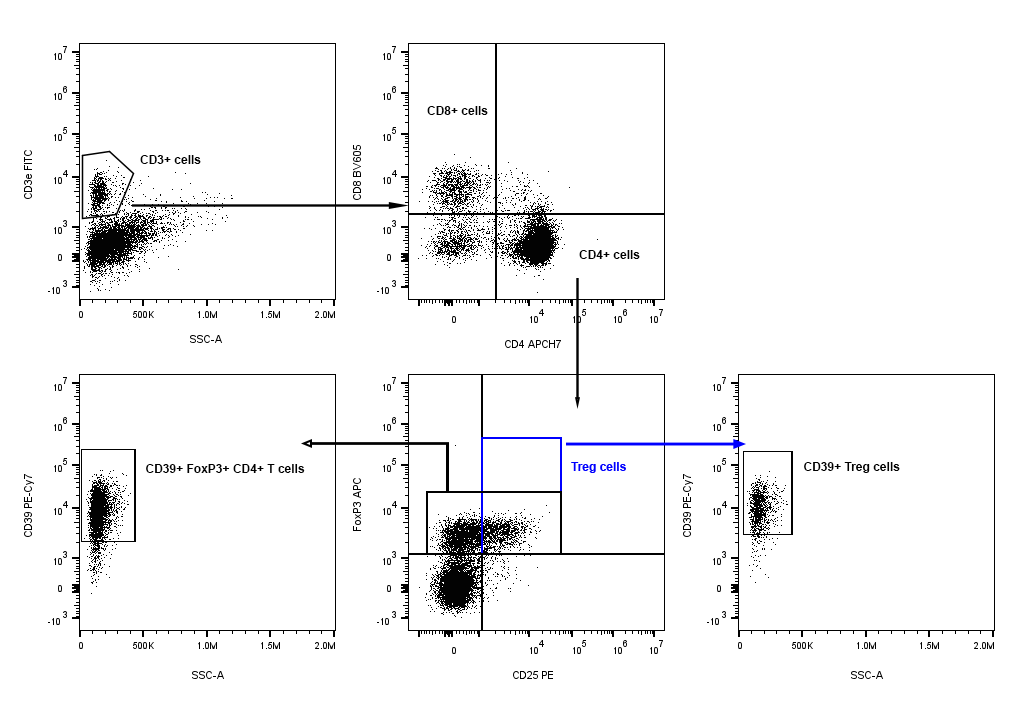

**Figure S16. Flow cytometry gating strategy for regulatory T cell (Treg) analysis.** Viable cells were first identified by exclusion of staining with a fixable viability dye (as described in *Figure S13*). Subsequent gating steps were applied to define CD4⁺ and CD8⁺ T cell populations, identify Treg cells, and assess CD39 expression within the Treg and FOXP3⁺ CD4⁺ T cell subsets.

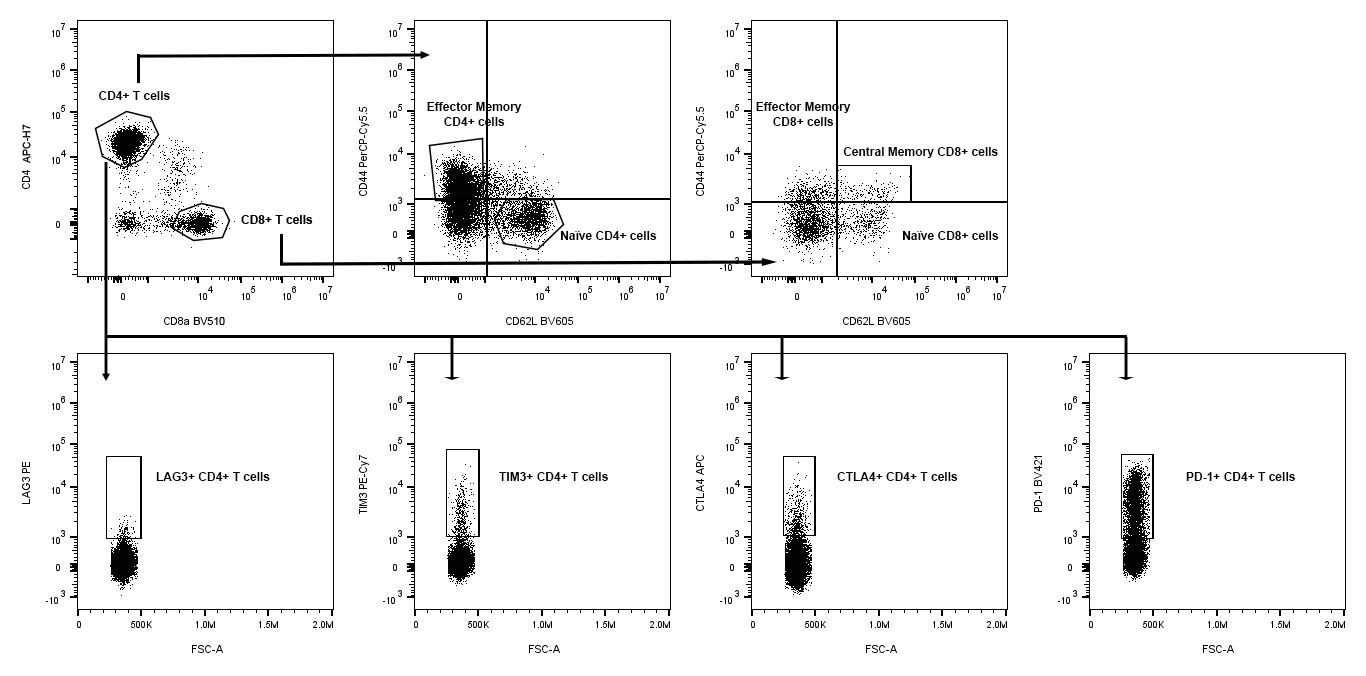

**Figure S17. Flow cytometry gating strategy for CD4⁺ and CD8⁺ T cell subpopulations and immune checkpoint analysis in CD4⁺ T cells.** Viable cells were first identified by exclusion of staining with a fixable viability dye (as described in *Figure S13*). Subsequent gating steps were applied to define CD4⁺ and CD8⁺ T cell populations, and to distinguish effector memory, central memory, and naïve subsets within each. Expression of immune checkpoint molecules (LAG-3, TIM-3, CTLA-4, and PD-1) was specifically assessed within the CD4⁺ T cell population.
